# Effect of Cholesterol on Membrane Partitioning Dynamics of Hepatitis A Virus-2B peptide

**DOI:** 10.1101/2020.12.28.424541

**Authors:** Samapan Sikdar, Manidipa Banerjee, Satyavani Vemparala

## Abstract

Understanding the viral peptide detection, partitioning and subsequent host membrane composition-based response is required for gaining insights into viral mechanism. Here, we probe the crucial role of presence of membrane lipid packing defects, depending on the membrane composition, in allowing the viral peptide belonging to C-terminal Hepatitis A Virus-2B (HAV-2B) to detect, attach and subsequently partition into the host cell membrane mimics. We conclusively show that the hydrophobic residues in the viral peptide detect the transiently present lipid packing defects, insert themselves into such defects, form anchor points and facilitate the partitioning of the peptide. We also show that the presence of cholesterol significantly alters such lipid packing defects, both in size and in number, thus mitigating the partitioning of the membrane active viral peptide into cholesterol-rich membranes. These results show differential ways in which presence and absence of cholesterol can alter the permeability of the host membranes to the membrane active viral peptide component of HAV-2B virus, via lipid packing defects, and can possibly be a part of general membrane detection mechanism for the viroporin class of viruses.

## 1. INTRODUCTION

Viruses pose a constant threat to human health. More so due to several viral outbreaks over the past decade, like Severe Acute Respiratory Syndrome (SARS), Middle East Respiratory Syndrome (MERS), Ebola and the recent worldwide pandemic caused by novel Coronavirus (nCoV/SARS-CoV2). The current situation necessitates not only detailed understanding of virus structure, but also how it interacts and subsequently partitions into host cell membranes.

Several studies [1–5] indicate that viruses encode small proteins/peptides that alter host membrane permeability, leading to enhancement of viral propagation. These membrane active viral components are usually hydrophobic / amphipathic in nature, typically unstructured and short with 60-120 amino acids in length [1, 2, 4]. Following membrane insertion, these components oligomerize to form pores/channels facilitating diffusion of small solutes and/or ions across the bilayer. Owing to this membrane permeation mechanism that embodies a channel-pore dualism[6], they are also known as viroporins. Viroporins are classified on the basis of number of transmembrane (TM) helices, which varies from one to three; and are further subdivided according to their membrane topology [1]. Class I viroporins include Influenza A Virus (IAV) Matrix 2 protein (M2) [3, 7] and Human Immunodeficiency Virus (HIV-1) Viral Protein Unique (Vpu) [8], which contain one transmembrane helix; while Class II and Class III viroporins like Hepatitis C Virus (HCV) protein 7 (p7) [8, 9], Poliovirus 1 (PV1) 2B protein and Coronavirus protein 3A [1, 2], contain multiple transmembrane helices. Several biophysical and biochemical techniques have provided useful insights into the structure and function of viroporins. [1–5] Owing to the small size and sensitivity to hydrophobic environment, structure determination based on biophysical techniques like solution and solid-state NMR, X-ray crystallography, electron microscopy have been limited to only IAV M2 [3, 7], HIV Vpu [8] and HCV p7 [8, 9] viroporins. Alternately, the PV1-2B [1, 2] has been widely studied through biochemical assays revealing a TM alpha-helical hairpin motif with cytosolic orientation of its N- and C-termini, which undergoes homo-oligomerization to form tetrameric transmembrane pores facilitating passage to small solutes and ions. The 2B protein is common in *Picornaviridae* family, which includes PV1, Foot-and-Mouth Disease Virus, and Hepatitis A Virus (HAV), to name a few. [1, 2]

Recently, a putative membrane interacting domain of 60 residues located at the C-terminal region of HAV-2B protein has been identified [10], the sequence of which is provided in SI, Fig S1. The 2B protein of HAV, like that of analogues from other picornaviruses, plays an essential role in membrane reorganization [11] and viral replication [12], but does not participate in calcium homeostasis or host membrane trafficking [13]. HAV-2B is also unusually longer (251 amino acids) compared to other picornavirus 2B proteins, that are typically, only around 70-110 residues in length. Shukla *et al.* [10] demonstrated the membrane permeabilizing property of the C-terminal region of HAV-2B based on biochemical assays. A synthetic construct of the 60-amino acid region adopted an alpha helical conformation under membrane mimicking conditions, as indicated by circular dichroism measurements, in agreement to reported membrane interacting domain of PV1-2B peptide [14]. Using membrane disruption assay, fluorescence spectroscopy and dynamic light scattering measurements, the authors showed pore forming ability of such peptide. In addition, the permeabilizing property of the peptide was observed to be dependent crucially on the lipid type and composition, indicating differential peptide activity against membranes of different cellular organelles. For instance, while endoplasmic reticulum membrane was relatively more prone to permeation, plasma membrane mimicking membranes, with significant cholesterol content, was least affected by the HAV-2B peptide. The authors reported that high cholesterol content impaired the membrane active property of HAV-2B peptide, although the mechanism of this remains unclear.

Although the crystal structure [15] of the N-terminal domain of HAV-2B has been determined in a different study, structural information pertaining to the putative membrane interacting region in the C-terminus is still lacking. In the absence of an experimentally determined structure, Shukla *et al.* [10] utilized molecular modelling methods to predict an alpha helical hairpin conformation for the HAV-2B peptide, a common structural fold reported for 2B peptides from *picornaviridae* family members. The authors performed short time-scale (~ 50-100 ns) molecular dynamics (MD) simulations of HAV-2B peptide in water and hydrated POPC membrane. All-atom simulations of multiple peptides in water revealed a preferred dimeric state. Based on additional simulations, in presence of POPC bilayer, the authors proposed that HAV-2B dimer interacts with membranes in a preferred orientation, characterized by terminals of both peptides making close contacts with lipid headgroups, while the alpha-helical hairpin motif remains exposed to solvent. This is in contrast to reported tetrameric arrangement of PV1-2B peptides in membrane, evident from biochemical experiments [16, 17] and MD simulation [18]. Similar orientational preference was also noted for monomeric HAV-2B peptide on membrane.

A majority of existing simulation studies on viroporins are related to membrane embedded states focusing on their preferred oligomeric arrangements [6, 18–21], channel/pore activity [22–25] and drug binding [26–28], to name a few. On other hand, atomistic and coarsegrained MD simulations of antimicrobial peptides [29–36] and fusion peptides [37–43] of enveloped-virus origin have explored peptide entry through partitioning and insertion into membranes. These studies provide detailed insight into effect of lipid type and composition on peptide-membrane modes of interaction and subsequent membrane reorganization by these membrane active peptides. However, simulations elucidating membrane permeabilization by non-enveloped virus proteins have been limited to HAV-2B [10], HAV-VP4 [44, 45] and gamma peptide of Flock House Virus (FHV) [46, 47]. These studies show peptide association with membrane, but do not capture membrane response upon such interaction and as such the mechanisms of permeation are less understood. Although several viroporins have been studied experimentally, the molecular basis of membrane destabilization caused by them remains elusive. Since viroporins are critical for propagation of viruses, and have been shown to be drug targets, it is essential that the molecular mechanism of their interaction with membranes be elucidated [26–28, 48, 49]. In addition, it is necessary to understand the role played by the membrane, its dynamic fluctuating state and how its composition facilitates entry of such viral peptides.

In this work, we have made efforts to gain microscopic insight into membrane response elicited by interactions of the HAV-2B peptide, which in turn facilitate their partition. We also investigate how presence of cholesterol mitigates the membrane disruption by the peptide. The amount of cholesterol is specific to membrane type and varies between 50% in mammalian plasma membrane to 10% in mitochondrial membrane, while other cellular organelles like Golgi bodies, endoplasmic reticulum have intermediate cholesterol content[10]. So, in our study we consider moderate cholesterol content for mixed bilayer simulations. Towards this goal, we perform extensive all-atom molecular dynamics simulations of HAV-2B peptide in (i) water, (ii) POPC and (iii) mixed POPC-Cholesterol (20%) bilayers. The conformational space sampled by the peptide predominantly depends on the environment. The peptide exhibits an extended conformation, with significant partitioning, in POPC bilayer; while it tends to adopt a compact conformation, without much partitioning, in the presence of cholesterol embedded bilayer. The HAV-2B peptide, partitioned into the POPC bilayer, acquires facial amphiphilicity via local segregation of its hydrophobic and hydrophilic residues along the peptide backbone. Our simulations strongly show that the peptide is capable of sensing membrane topography in the form of lipid packing defects and that its partitioning dynamics is primarily driven by discrete events of hydrophobic residue insertions into such packing defects. The peptide insertion further destabilizes the membrane by inducing global thinning and enhanced mobility of lipid tails. However, such membrane induced partitioning dynamics is considerably inhibited in cholesterol rich membrane patches due to the presence of smaller and fewer packing defects. The present simulation study, to the best of our knowledge, provides first atomistic insight into membrane partitioning dynamics of non-enveloped viral peptides characterized by sensing and stabilizing of lipid packing defects and a direct evidence for their role in membrane remodelling.

## 2. RESULTS

In our attempt to understand the effect of surrounding medium on conformational dynamics of HAV-2B, we performed extensive simulations of the peptide in presence of aqueous solution, pure POPC bilayer and mixed bilayer with POPC and Cholesterol (20%), the corresponding system details being provided in SI Table S1. The modelled peptide[10] served as the initial configuration for peptide-water systems. We calculated root mean squared deviation (RMSD) per residue considering backbone C-α atoms averaged over last 50 ns (see SI, Fig S1), with the initial modelled peptide serving as the reference. In aqueous medium the peptide undergoes significant deviation from the initial modelled configuration (Fig S1A), especially at the terminal residues, suggesting the rich conformational space explored by the peptide in solution phase. The ensemble averaged structure of HAV-2B peptide in water is shown in Fig S1B. The bilayer simulations are started using a representative peptide configuration from the equilibrated aqueous solution ensemble. In presence of the hydrated POPC bilayer, large scale deviation from initial solution state is observed, indicating conformational re-arrangement of the peptide upon going from solution to membrane phase (Fig S1A). However, the peptide in presence of cholesterol, exhibits residue level average deviations largely similar to that observed in aqueous medium (Fig S1A). Owing to these observed differences, we first present a detailed account of conformational heterogeneity of HAV-2B peptide induced by nature of surrounding medium: (i) water, (ii) POPC bilayer and (iiii) POPC-Chol(20%) bilayer. Further, we elucidate the peptide’s mode of interaction with membrane and role in modulating membrane properties. In order to study the latter, we perform additional simulations of pure POPC bilayer and POPC+20% cholesterol, serving as control systems.

### 2.1. Peptide conformation

#### (a) in Water

The conformational states of HAV-2B peptide are differentiated based on average radius of gyration (Rg). The Rg is computed as the average distance of backbone heavy atoms from their centre of mass, the distribution, P(Rg), (Fig S2A), being generated from equilibrated part (see SI Table S1) of the trajectories and provides a measure of molecular dimension. The HAV-2B peptide in aqueous solution exists in a multitude of conformations ranging from somewhat compact (~10 Å) to extended-like states (~15 Å).

The peptide conformations are further characterized by monitoring the presence of different secondary structural elements, assigned using the STRIDE [50] algorithm implemented in VMD [51]. The secondary structure percentage (SS %) of each residue calculated over the equilibrated trajectories represents the population of different SS-elements explored by each residue during the course of simulation. The residue based SS % of HAV-2B peptide in the solution phase is shown in SI (Fig S3A). The N-terminal, V1-G16 and C-terminal, A43-L60 residues predominantly exhibit “turn” or “random coil” like conformations, typical of an unstructured dynamic loop, as also evident from the average structure illustrated in Fig S1B. I17 to L25 and I33 to Y42 form two well defined alpha-helices of the hairpin motif (see Fig S1B). These helices are connected via an unstructured loop (N26-E29) followed by a residue triad H30-S31-H32, which undergoes frequent transitions between “turn”, 3_10_-helix and alpha-helix.

Inter-helical crossing angle (Ω) also serves as a suitable conformational parameter to describe alpha helical hairpin motifs [52]. In order to probe Ω, we define two axes corresponding to the two helices of this motif and calculate the crossing angle between them. The vector between C-alpha atoms of L18 and I22 represents the first helical axis *(hP),* while that of L36 and M40 represents the second *(h2)* (see Fig S1B). The equilibrium distribution, P(Ω), shown in SI Fig S2B indicates a broad spectrum of accessible inter-helical angles varying between 120°-180° in water.

#### (b) at POPC-Water interface

The HAV-2B peptide explores more extended conformations, compared to its solution phase structure, with Rg varying predominantly between 14-20 Å, in vicinity of POPC bilayer as indicated by P(Rg) (SI, Fig S2A). The presence of the long stretched C-terminal tail as illustrated in Fig 1A contributes to the observed large Rg in POPC bilayer. The peptide likely gets locked into this extended conformational state upon interaction with POPC bilayer as discussed in Section 2.2. The SS % (Fig S3B) indicates loss of alpha-helical propensity for a few polar residues. The residues N41 and Y42 located at end of the second helix completely lose their helical profile (see Fig 1A). The observed loss in helical nature of these polar residues may have implications in partitioning dynamics as discussed in details in Section 2.2. Barring these residues, the secondary structure of HAV-2B peptide at the hydrated POPC interface remains largely similar to that in solution phase. In contrast to aqueous solution, the POPC bilayer interface induces reduced fluctuations in inter-helical angles, Ω, leading to tapered distributions (SI Fig S2B), with peak around 165° resembling an anti-parallel helical conformation characteristic of a hairpin structure. Such values are in close agreement to reports on inter-helical angles of helix packing motifs in membrane proteins[53].

**Fig 1.**
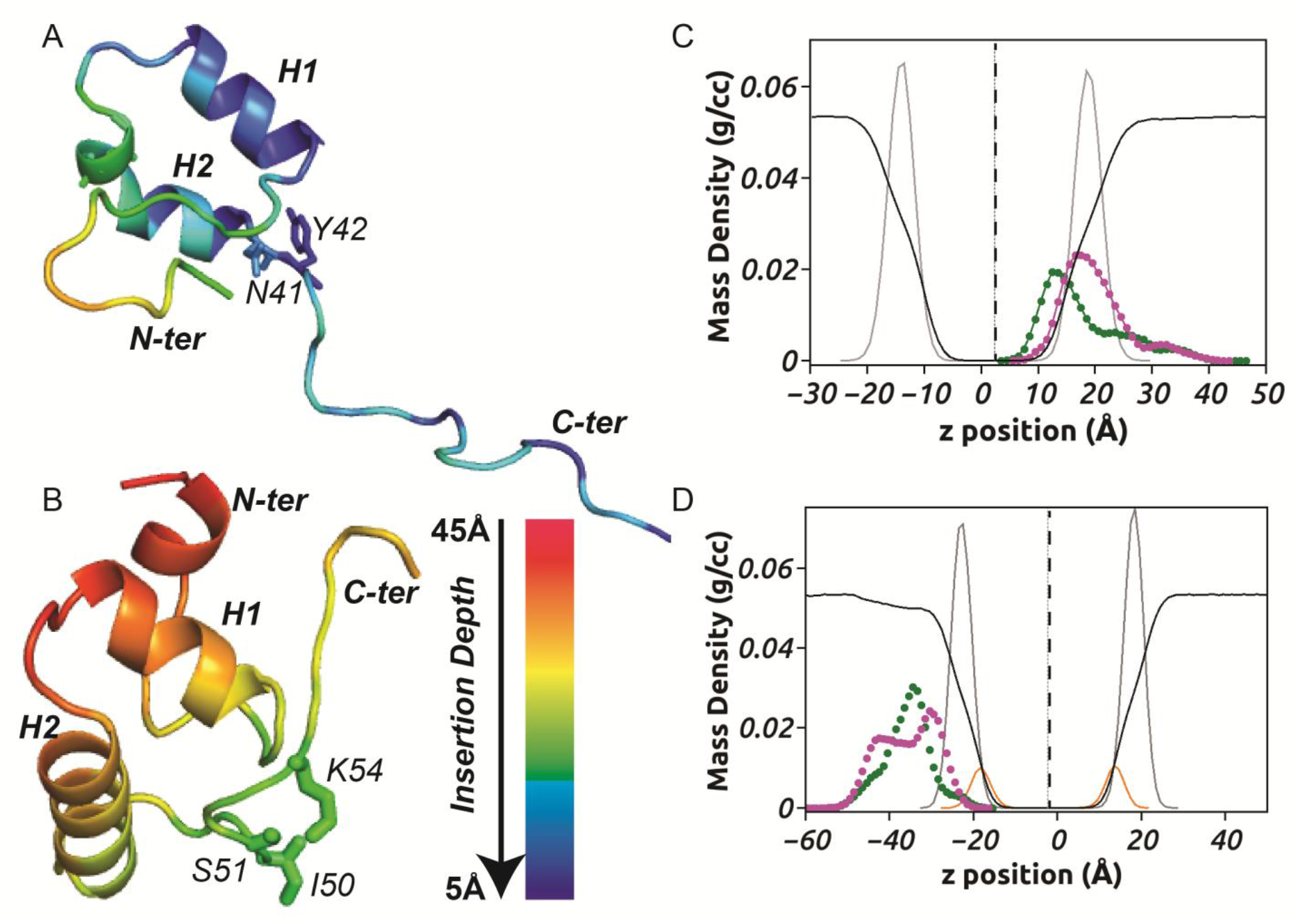
Conformational differences of peptide in presence or absence of cholesterol affecting peptide partitioning into membrane. The conformational difference of HAV-2B peptide indicating (A) an extended state in POPC bilayer and (B) a compact state in POPC-Cholesterol (20%) bilayer. The H1 and H2 helices form the alpha helical hairpin motif. The average distance of peptide residues from respective bilayer centres are indicated through the colour scale, representing insertion depths. It is to be noted that lower the value, the deeper is the insertion, indicated in blue while red indicates shallow or absence of insertion. The peptide residues N41 and Y42 lose their helical conformation owing to deep partitioning in POPC bilayer. In presence of cholesterol, I50, S51 and K54 from C-terminal tail are more close to bilayer centre than rest of the peptide. (C) Mass density profile along the bilayer normal (z-direction) of HAV-2B peptide showing segregation of hydrophobic and hydrophilic residues towards bilayer centre (black dotted line) and POPC headgroup-water interface, respectively. (D) The z-component of mass density profile of HAV-2B peptide in POPC-Cholesterol (20%) bilayer showing overlapping densities of hydrophobic and hydrophilic residues near the solvent proximal interface. The hydrophobic and hydrophilic components are shown in green and magenta, respectively. The density profiles of POPC headgroups, cholesterol and water (reduced by a factor of 10) are shown in gray, yellow and black, respectively.

#### (c) at hydrated POPC-Chol(20%) interface

In contrast to the predominantly extended conformations in POPC bilayer, the peptide near the POPC-Chol phase is found to be in a compact state, even more than the sampling in the solution phase, as can be seen by the narrow and peaked distribution of P(Rg) around 11 Å (Fig S2A). This compact conformation of the HAV-2B peptide in POPC-Chol(20%) bilayer, shown in Fig 1B is largely due to the N- and C-terminal tails which fold back towards the helices unlike adopting the stretched conformation as in POPC bilayer. The distribution of inter-helical angle (Fig S2B) with mean value of Ω close to 150° indicates that the two helices are far from being anti-parallel leading to a distorted hairpin motif, also evident from Fig 1B. Further, the hairpin conformation and few N-terminal residues are characterized by an increased helical content (Fig S3C), which is significantly different from that in POPC bilayer.

### 2.2 Peptide-Membrane mode of interaction

Besides the conformation, the orientation of the peptide relative to membrane surface also governs its mode of interaction with the lipids. Several studies highlight the importance of orientational preference [30, 35, 54–56] of membrane-active peptides^30, 35, 66-68^ related to membrane permeabilization mechanisms. In this study, the peptide orientation is described by the hairpin tilt angle (τ) with respect to the bilayer normal. Since the two helices of the hairpin are nearly anti-parallel at the membrane-water interface, it is sufficient to consider one of the helical axes. Accordingly, τ is defined as the angle between the first helix axis and the membrane normal as illustrated in SI, Fig S2C. The orientation dynamics of the peptide is probed through equilibrium fluctuations in τ (SI, Fig S2D). The HAV-2B peptide adopts a near-horizontal orientation with <τ> ~ 70° parallel to the POPC bilayer in absence of cholesterol. The near-horizontal orientation ensures maximum interaction of the alpha-helical hairpin motif with the POPC bilayer owing to increased surface area of contacts. In contrast, the H1 helix of the peptide orients more along the normal of POPC-Chol(20%) bilayer while, the other helix adopts an inclined orientation due to the distorted hairpin structure in vicinity of cholesterol embedded bilayer. Such an orientation in presence of cholesterol significantly reduces the contact surface of the peptide with the bilayer. The orientational difference thus translates to change in residue contacts which can have significant effects on peptide-membrane interactions as indicated in the following section.

To investigate the nature of the peptide interactions with the membrane in absence and presence of cholesterol, we determine residue contact probabilities (see SI Table S2) from last 50 ns trajectories, representing the membrane-embedded/adsorbed state of the peptide. In absence of cholesterol, a majority of HAV-2B peptide residues have contact probabilities greater than 75%. These residues, both polar and hydrophobic in nature, belong to the hairpin motif and the C-terminal tail. They form extensive contacts owing to the horizontal alignment of the alpha-helical hairpin motif and the stretched C-terminal tail along the POPC membrane. This in particular ensures that the peptide gets locked in the extended conformation upon interaction with POPC bilayer. However, due to the orientational and the conformational preferences in cholesterol mixed bilayer, the surface contacts are limited to I50, S51 and K54 from the C-terminal tail.

The peptide localization is quantified through its insertion depth into the membrane milieu. The depth of insertion is calculated as the distance of z-component of centre of mass (z C.O.M) of each peptide residue from the bilayer centre along the membrane normal. The smaller the value of insertion depth, the deeper is the insertion. Fig 1A shows the peptide with colour coded residues representing their relative distance from centre of the POPC membrane. The N-terminal tail is far (~20 Å) from membrane interior, while the alpha-helical hairpin motif and the long stretched C-terminal tail are located in close proximity (~5-10 Å) to bilayer centre indicating deep insertion, resulting in extensive contact formation with lipid molecules. It is interesting to note that the polar residues N41 and Y42 (see Fig 1A) partition deep into the membrane and thus exhibit high contact probability (SI, Table S2). Their backbones reside deep into the hydrophobic membrane interior while their polar side-chains face out towards the hydrophilic POPC headgroups. Such an arrangement accounts for the observed loss of backbone helical structure as described earlier. However, in presence of cholesterol, the membrane interior remains inaccessible to majority of the peptide residues as indicated by the reduced insertion depth in Fig 1B. The hairpin motif along with both the termini remains far from membrane interior leading to reduced contacts with the membrane. Only the residues, I50, S51 and K54 are observed to be in close proximity to POPC-Chol(20%) bilayer.

Peptide insertion is often driven by membrane induced partitioning, widely reported for antimicrobial peptides [30, 34, 55], polymers [31–33, 35, 36, 57] and other membrane active molecules [58–60]. We confirm this by computing equilibrium density profiles of peptide and membrane components along the bilayer normal (z-axis). The z-density profile of HAV-2B peptide in POPC (Fig 1C) and POPC-Chol(20%) (Fig 1D) bilayer reflects the distribution and localization of the hydrophobic and hydrophilic residues along the z-axis. That most residues of the HAV-2B peptide undergo deep insertion in POPC bilayer, is also reflected in Fig 1C. Significant density contribution by hydrophobic residues is observed to reside below lipid headgroups, tailing towards the membrane centre. The density peaks in Fig 1C indicate segregation of hydrophobic and hydrophilic moieties of the peptide towards bilayer centre and headgroups, respectively, leading to membrane induced partition. The density profiles extend towards the solvent proximal interface due to freely fluctuating N-terminus of HAV-2B peptide being exposed to hydrating water (Fig 1C). These results clearly indicate that the hydrophilic residues of the peptide favour localization near the polar interface and facilitate initial contacts with membrane surface, while the hydrophobic ones prefer membrane interior tending to lower the insertion depth, suggesting a strong facially amphiphilic conformation of the peptide in POPC membrane. A representative snapshot of HAV-2B peptide embedded in POPC bilayer is illustrated in Fig 2A. The side-view shows deep insertion of the alpha-helical hairpin motif and the C-terminal tail as a result of partitioning between hydrophobic and hydrophilic residues indicated in green and magenta, respectively. A 90°-rotated top view shows the extended conformation of the peptide localized on the membrane x-y plane. However, in presence of cholesterol, the density profile of the peptide rarely overlaps with that of lipid headgroups (Fig 1D) substantiating our observation that only few residues are able to make surface contacts with the lower leaflet of POPC-Chol(20%) bilayer and that the membrane interior remains largely inaccessible to the peptide. Moreover the density profiles of hydrophobic and hydrophilic groups of the peptide are overlapping with peak positions in solvent. This clearly demonstrates that the presence of cholesterol precludes partitioning of the peptide leading to superficial adsorption on membrane surface. Our results are in excellent agreement to existing experimental observations based on membrane disruption assay, which indicated artificial membranes with high cholesterol content impaired the permeation property of HAV-2B peptide.[10] A representative of the membrane adsorbed state of HAV-2B peptide on POPC-Chol(20%) bilayer is illustrated in Fig 2B. The alpha-helical hairpin motif and the N-terminal tail remain solvent exposed while a handful of residues belonging to the C-terminal tail form stable contacts with lipid headgroups (see SI Table S2). The top view indicates a more “confined” localization on POPC-Chol(20%) membrane surface due to a compact globular conformation. Owing to lack of interaction with the membrane, the peptide tends to adopt a compact conformation under the effect of enhanced intra-peptide interactions.

**Fig 2.**
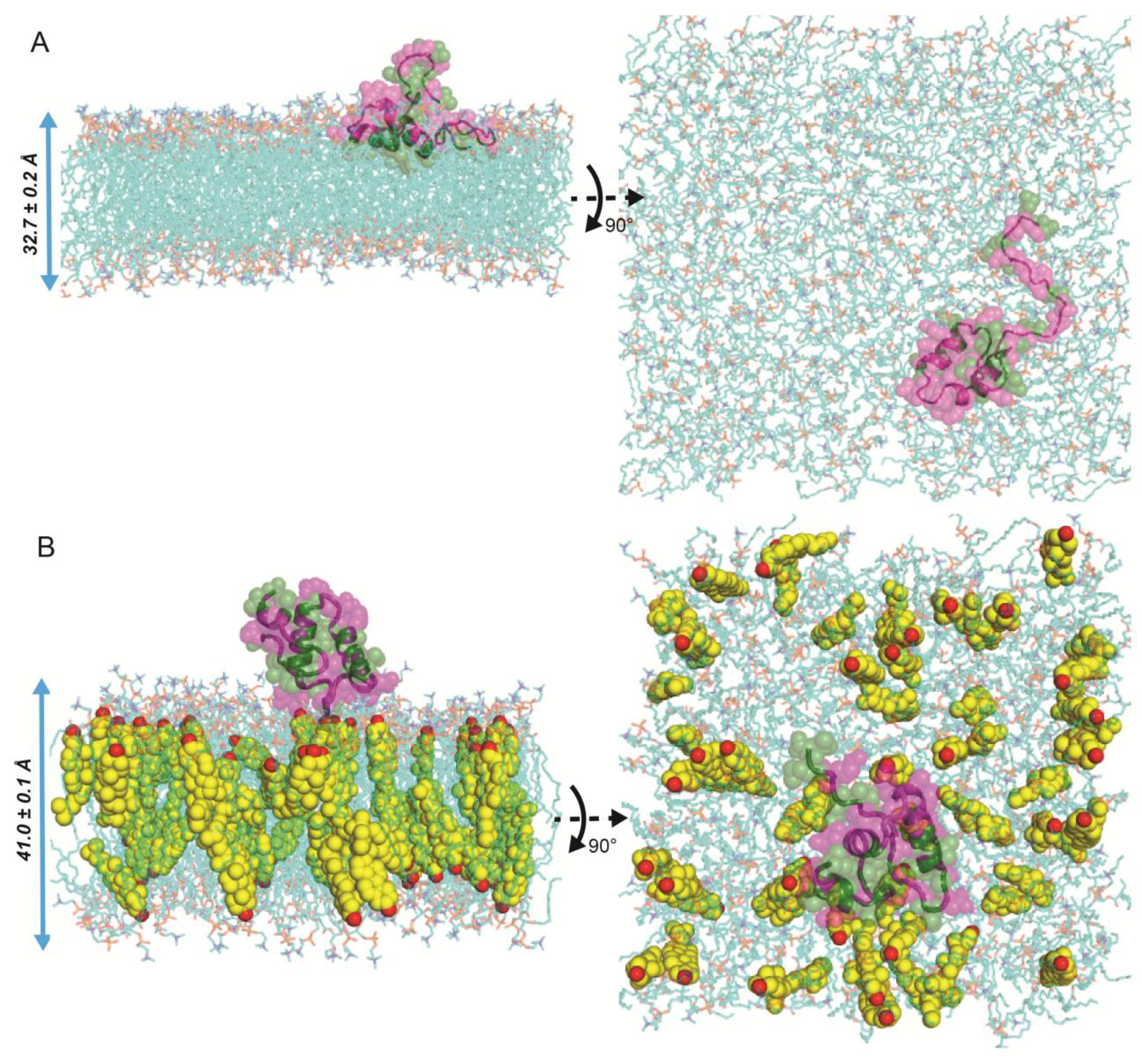
Representative snapshots (side- and top-view) of HAV-2B peptide localization in membrane. (A) Membrane induced peptide partitioning in POPC bilayer and (B) surface adsorption of the peptide on POPC-Chol(20%) bilayer. In the snapshots, the POPC molecules are indicated by transparent sticks (cyan), cholesterol molecules by spheres (yellow) and peptide by cartoon and transparent spheres with hydrophobic residues in green and hydrophilic residues in magenta. Water molecules are not shown for clarity. The bilayer thickness is indicated by blue arrow.

### 2.3 Membrane remodelling

Partitioned membrane active agents like peptides and polymers have been shown to induce significant changes in the membrane state or induce membrane remodelling [5, 30, 33, 35, 36, 58, 59, 61]. In this section, we investigate if a similar modulation of membrane organization is induced by the peptide interactions and the role played by presence of cholesterol. The two peptide-free systems, pure POPC and mixed POPC-Chol(20%) bilayers act as controls to compare peptide induced changes in membrane properties, like bilayer thickness, surface area/lipid and lipid acyl tail ordering. We first draw comparison between the two control systems to understand the role of cholesterol and then compare the respective controls with the peptide based systems to understand the effect of peptide partitioning on membrane properties.

#### (a) Bilayer thickness

We compute 2d-thickness profiles with a 2 Å x 2 Å grid resolution along xy-plane corresponding to final snapshots of control and peptide-membrane systems (see Fig 3). The lateral thickness profile of control POPC bilayer (Fig 3A) is moderately homogeneous with local thickness varying between 35-45 Å. The addition of cholesterol to the control pure POPC system leads to membrane thickening but the corresponding 2d-thickness profile (Fig 3D) is comparatively uniform unlike pure POPC system. This is also evident from the ensemble averaged values (SI, Fig S4A) indicating enhanced thickness of ~ 43 Å in POPC-Chol(20%) bilayer compared to 39 Å in pure POPC, consistent with existing experimental[62] and simulation[63–65] studies.

**Fig 3.**
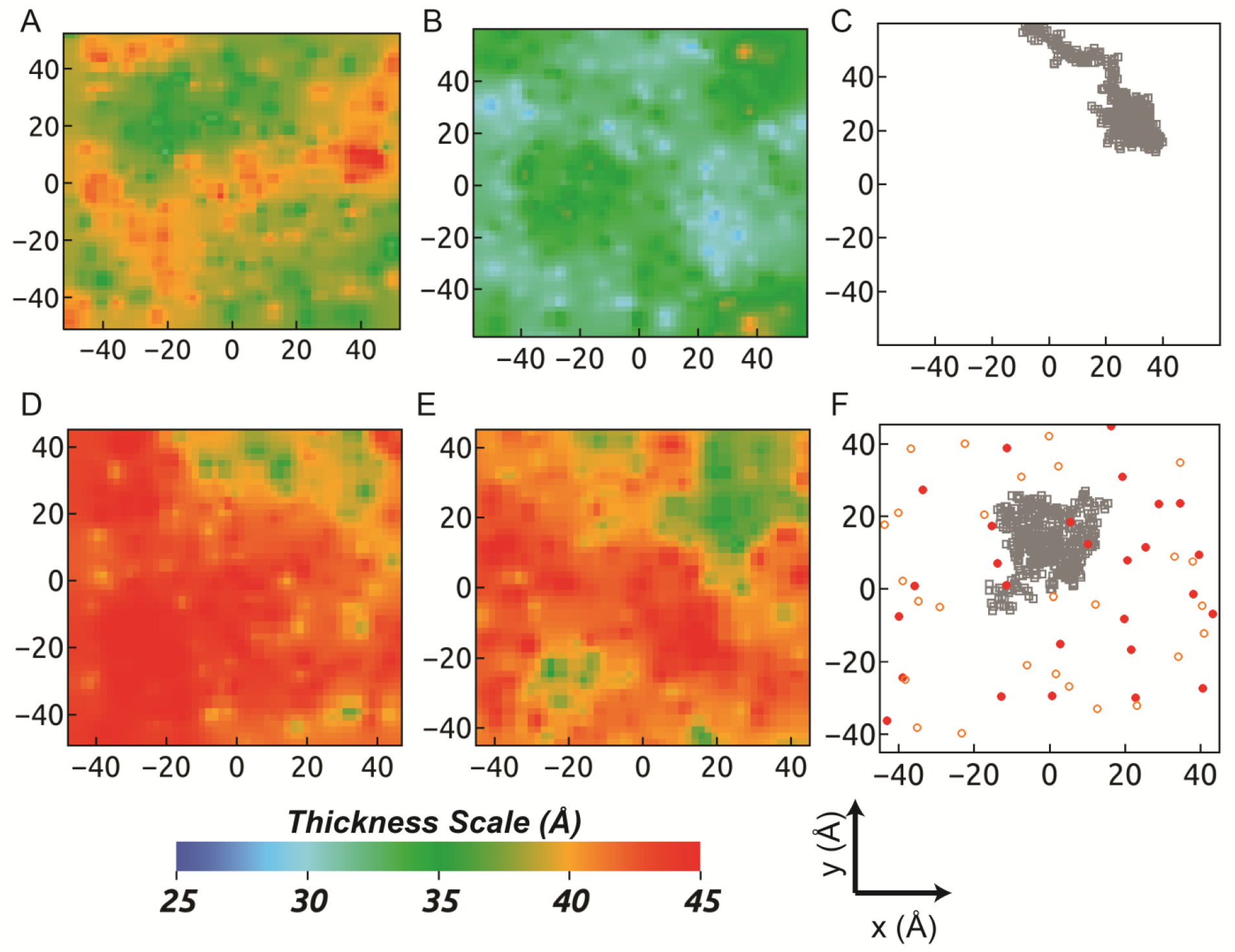
2-D grid based membrane thickness in x-y plane. The thickness maps are generated using a 2Å resolution along each direction for representative snapshots from equilibrated trajectories of systems considered in the present study. The membrane thickness of (A) pure POPC bilayer is significantly reduced upon interaction with (B) HAV-2B peptide. The peptide localization on membrane plane is shown in (C). Increased membrane thickness is observed in (D) mixed POPC-Chol(20%) and (E) HAV-2B POPC-Chol (20%) bilayers. (F) The peptide (gray) is localized on the lower leaflet. The cholesterol headgroups in both leaflets are shown in red filled circles (top) and orange open circles (bottom). The cholesterol-rich regions exhibit more thickness.

The peptide partitioning into the POPC bilayer leads to a more uniform distribution of local thickness as evident from Fig 3B, but significantly lowered (~ 30-35 Å) compared to the respective control. It is interesting to note that the membrane thinning effect is induced globally, albeit more pronounced in vicinity of the peptide (Fig 3C). The ensemble averaged global bilayer thickness values (see SI, Fig S4A) reflect ~ 15% reduction upon peptide insertion. However, superficial adsorption of HAV-2B peptide on POPC-Chol(20%) bilayer surface has only “local” thinning effect (Fig 3E-F) resulting in just 4% lowering of overall membrane thickness compared to the control cholesterol mixed bilayer. Further, the presence of peptide induces inhomogeneity in the 2d-profile characterized by local thickness varying between 35 Å close to its location and 45 Å in cholesterol rich areas.

#### (b) Average Surface area per lipid

The change in bilayer thickness is known to be correlated to change in surface area per lipid (SA/lipid), the values of which averaged over the equilibrated trajectories are shown in SI, Fig S4B. A comparison between the two controls reveal a significant decrease in SA/lipid of POPC molecules owing to the condensing effects of cholesterol, vastly reported in literature[59, 63–66]. However, peptide insertion enhances the SA/lipid considerably from 65.3 Å^2^ in pure POPC to 81.7 Å^2^ in HAV-2B-POPC system. As the peptide fails to partition into the POPC-Chol(20%) bilayer, only modest increase (~ 5%) in SA/lipid is observed, which can be explained as within the natural fluctuation of these values (see Fig S4B).

#### (c) Order Parameter

The lipid tail flexibility is intricately related to membrane thickness; for instance, the bilayer condensing effects of cholesterol results in a more ordered phase[66]. The lipid tail order is determined from deuterium order parameter (SCD) accessible from NMR experiments and MD simulation, calculated as S_CD_ = *½* | *<3 cos^2^ θ - 1>* |, *θ* being the angle between C-H bond and bilayer normal. Higher S_CD_ corresponds to reduced flexibility of lipid acyl chains and vice-versa. The order parameter, S_CD_, is computed over the equilibrated trajectories for both saturated (sn-1) and unsaturated (sn-2) acyl chain carbon atoms from the two leaflets (top and bottom) separately and shown in Fig 4. Upon comparison between the two controls, we observe the enhanced ordering of both sn-1 and sn-2 chains in presence of cholesterol as expected. The presence of cholesterol reduces the POPC acyl chain flexibility by intercalating in void spaces between them so as to reduce the surface area per lipid leading to bilayer condensation. Since membranes act as incompressible fluids, the condensing effect enhances the cholesterol mixed bilayer thickness.

**Fig 4.**
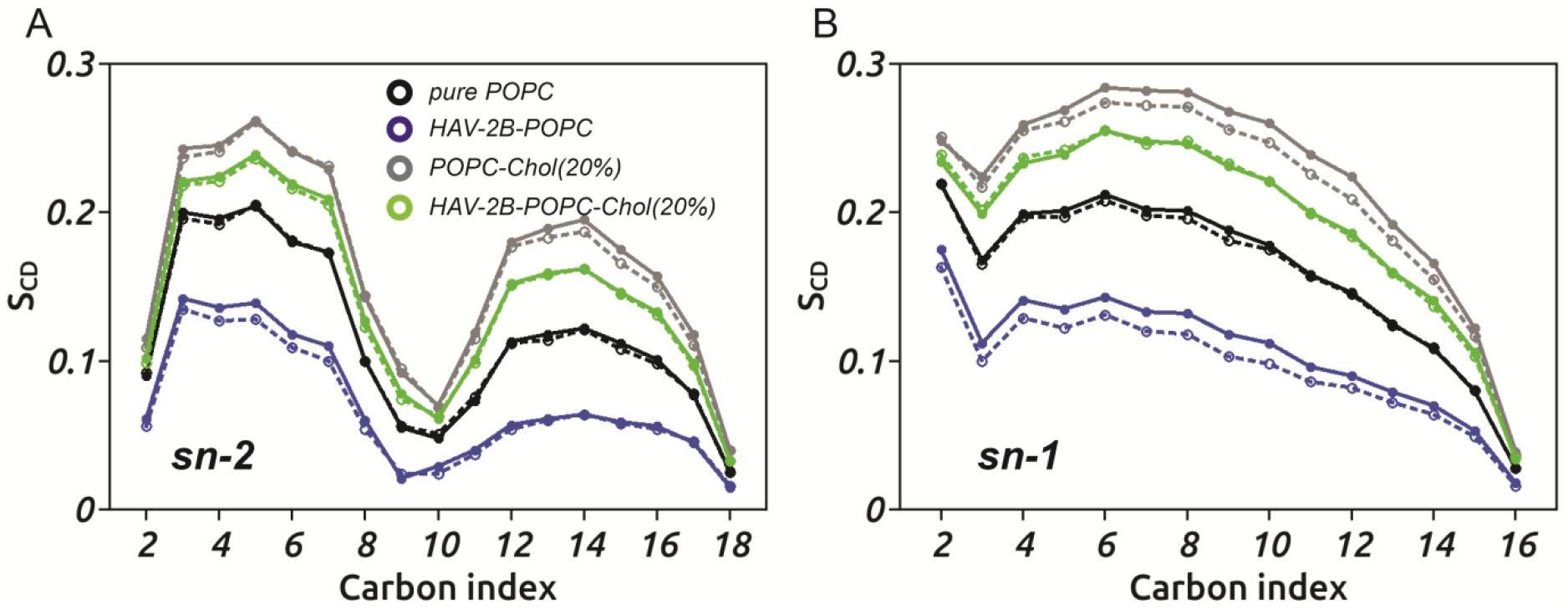
POPC lipid tail order parameters, S_CD_ corresponding to top (solid symbol and solid line) and bottom (open symbol and dotted line) leaflets for (A) sn-2 and (B) sn-1 chains.

In contrast, the peptide partitioning into the POPC bilayer results in strong membrane disorder, characterized by increased flexibility of POPC acyl tails. The enhanced flexibility of lipid tails in presence of peptide is further ascertained through distribution of both sn-1 and sn-2 chain tilt angles of POPC. To this end, the angle between the vector defined by the first and the last aliphatic carbon atoms of each chain and the membrane normal is computed and the ensemble averaged distribution for the same is shown in SI Fig S5. The distributions of sn-1 and sn-2 chain tilt angles indicate a decrease in peak height and simultaneous shift in peak position to higher angular values in presence of the HAV-2B peptide compared to the control POPC. Such broad distributions of tilt angles in HAV-2B-POPC system reflect enhanced flexibility of lipid acyl tails and complement the observed decrease in order parameters. The observed global thinning of POPC membrane in presence of HAV-2B peptide results in lateral expansion of bilayer. Such lateral expansion causes the increase in surface area of lipids as well as the flexibility of lipid tails. This reorganization of membrane properties influenced by deep partitioning of HAV-2B peptide contributes to understanding of the molecular basis of membrane destabilization and thus elucidates the membrane active property of this viral peptide consistent with experimental observations[10]. Moreover as the presence of cholesterol precludes peptide partitioning, the corresponding acyl chain S_CD_s show intermediate values between that of pure POPC and mixed POPC-Chol(20%) bilayer, indicative of mild disordering effects.

It is interesting to note that although the peptide partitions deep into the top leaflet of POPC bilayer, the induced disorder is reflected in both leaflets (see Fig 4 and S5) to a similar extent signifying inter-leaflet coupling. The inter-leaflet coupling may arise due to bilayer registry, where domain formation in a leaflet drives the same in the opposing leaflet.[67] Various factors like membrane remodelling, lipid compositional asymmetry, acyl chain interdigitation are known to regulate bilayer registry.[67, 68] The enhanced disorder of POPC acyl chains due to deep partitioning of the peptide facilitates strong interdigitation of lipid tails so as to elicit a coupled response in both leaflets. Such entropy driven coupling mechanism is known for liquid-disordered phases of single component symmetric bilayers.[69]

### 2.4 Role of Membrane Defects on peptide interaction

#### (a) Distribution of defect sizes

The presence of HAV-2B peptide modulates the membrane organization so as to induce lipid tail disordering to a large extent. Such perturbation of local arrangement of lipids in a bilayer gives rise to lipid packing defects, which in turn affects peptide-membrane interaction in a co-operative manner.[70, 71] These packing defects are characterized by low lipid density and exposure of the membrane hydrophobic core to hydrating solvent. In view of recent investigations[34, 36, 70–77] revealing the role of such lipid packing defects in driving membrane induced partitioning, we analyse the same for all systems considered in this study using Packmem software[78]. The underlying algorithm divides the x-y plane into 1 Å × 1 Å grids and scans along the z-direction from solvent interface upto 1 Å below the average level of C2 atoms of glycerol moities (see SI, Fig S6) of POPC to identify voids: regions of no lipid density. The packing defect sites are qualitatively characterized into “Deep” (below C2 atom) and “Shallow” (above C2 atom) and further quantified by size given by their area (A) of occupation. The calculation performed for each frame reveals the number, size and location of different defect sites, both “Deep” and “Shallow”. Summing up the individual areas of all “Deep” (“Shallow”) defect sites in a given frame and normalized by the area of leaflet defines the “Deep” (“Shallow”) defect area fraction (*f*) in the given frame. The probability distributions of defect sizes, P(A), are drawn from the last 250 ns data of all systems and fitted using single exponential decay functions, *Bexp(-A*/Π), considering A > 15 Å^2^ and P(A) ≥ 10^-4^, as suggested by the Packmem[78] authors. The decay constant, Π, characteristically defines the membrane topology in terms of interfacial defects. The higher the value of Π, the more abundant are large defect sites. The defect area fraction, *f*, paints an overall picture of extent of lipid packing defects in a given frame, while the defect size distribution provides a more detailed insight into abundance of defect sites of varying sizes accumulated over time.

We first compare the lipid packing defects of the two control systems: pure POPC and mixed POPC-Chol(20%) bilayers. We characterize the two controls based on the total area attributed to “Deep” defects per leaflet in a given frame. However, the two controls significantly vary in leaflet surface area due to the condensing effects of cholesterol. Hence, we compute the defect area fraction, *f*, normalized by the leaflet area of the respective systems. The corresponding distributions of the defect area fraction, P(*f*), is shown in Fig 5A. In pure POPC bilayer, the P(*f*) is broad indicating a significant fraction of leaflet area being covered by “Deep” defects. In presence of cholesterol, this total defect area shrinks as evident from the sharp distribution, P(*f*), with a peak shift towards lower values of defect area fraction. Now such decrease in defect area under the influence of cholesterol may arise due to limited occurrence of defect sites or reduction in the size of defect sites or a combination of both. In order to understand the underlying cause, we first calculate the number of “Deep” defect sites per leaflet (*nsites*) in each frame. The time evolution of *n_sites_* for the two controls is shown in Fig 5B. The number of “Deep” defect sites per leaflet is clearly reduced in presence of cholesterol indicating lower occurrence of such sites compared to that in pure POPC bilayer. This is illustrated through representative snapshots showing prevalence of different defect sites in the two controls (SI, Fig S7). We then estimate the relative population of defect sites of various sizes through the defect size distribution function, P(A) illustrated for pure POPC in Fig 5C and mixed POPC-Chol(20%) in Fig 5D, respectively. The P(A) is shown in semi-log scale, where inverse of the fitted slope yields the corresponding defect area constant, Π in Å^2^. The defect size distributions are similar in both systems with comparable Π, (see Table 1). The similarity is observed not only for “Deep” defects but also preserved for “Shallow” defects. Although, the number of defect sites is considerably reduced in cholesterol embedded bilayer, the relative population of different defect sizes remain almost similar to that of pure POPC bilayer. Hence, the observed lower value of defect area fraction is a direct consequence of low abundance of defect sites in cholesterol embedded bilayers rather than reduction in size of defects.

**Fig 5.**
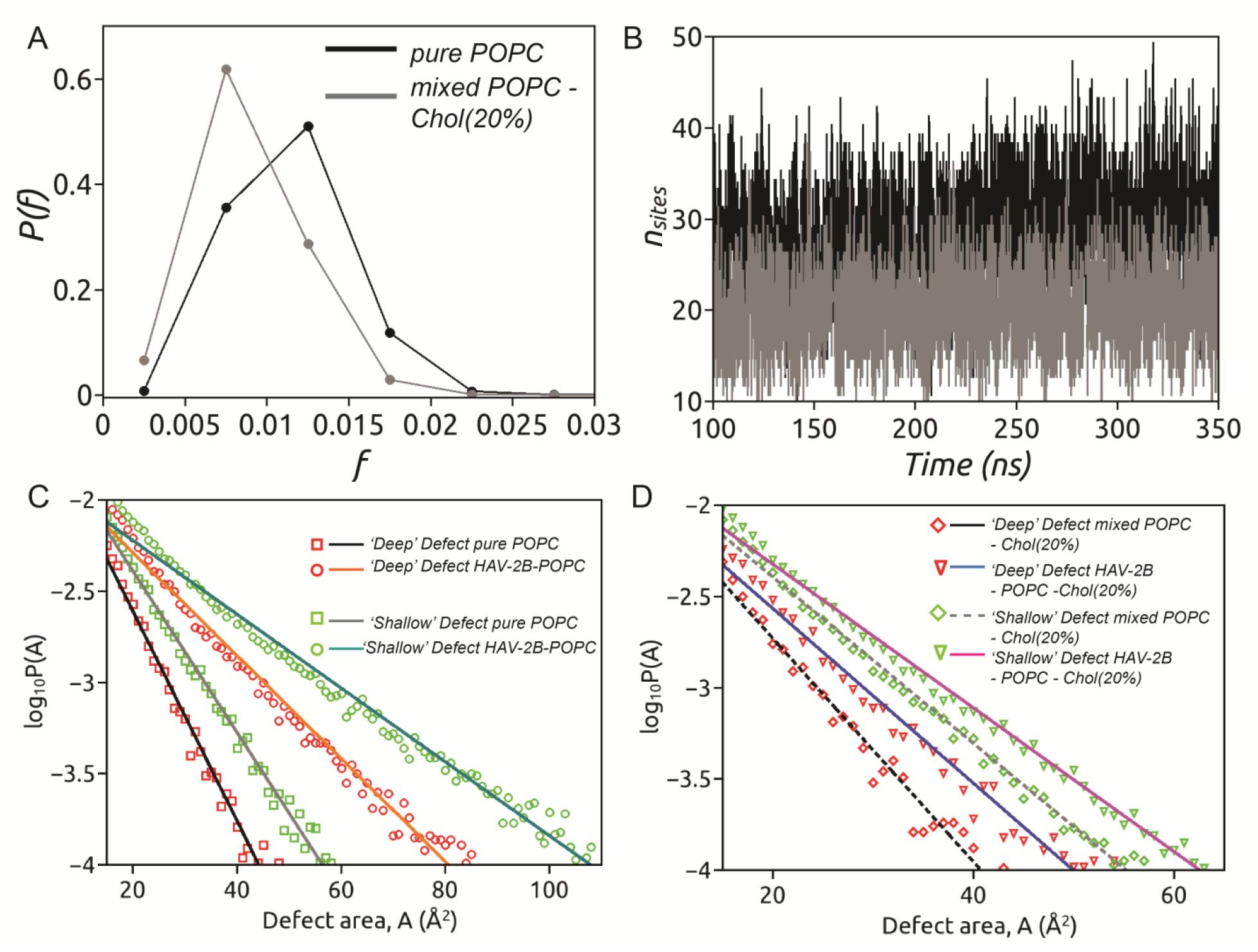
Quantification of lipid packing defects. (A) The distribution of defect area fraction, P(*f*) corresponding to “Deep” defects found in the two control systems: pure POPC and mixed POPC-Chol(20%). It provides a measure of the extent of defects in a given frame expressed as a fraction of the leaflet area. (B) The time evolution of number of “Deep” defect sites per leaflet *(n_sites_)* for the two controls. The effect of peptide partitioning on size distribution of lipid packing defects is illustrated for (C) HAV-2B-POPC and (D) HAV-2B-POPC-Chol(20) systems in comparison to the two controls. The probability distributions of defect sizes (in Å^2^), log_10_P(A), computed over last 2500 frames (i.e. last 250 ns), 100 ps apart considering both leaflets, are shown for ‘Deep’ (red) and ‘Shallow’ (green) packing defects. The lines represent single exponential fits considering defect sizes larger than 15 Å^2^ and log_10_P(A) ≥ 10^-4^ with corresponding area constants, π, listed in Table 1.

**Table 1.** Defect area constants, π (in Å^2^) derived from single exponential fits to packing size probability distributions as described earlier. The standard deviations are calculated using block averaging of each data set.

| **Systems** | **Deep defect**<br>**area constant,**<br>**$\pi$ ($\text{\AA}^2$)** | **Shallow**<br>**defect area**<br>**constant, $\pi$**<br>**($\text{\AA}^2$)** |
| --- | --- | --- |
| Pure POPC | $8.3 \pm 0.6$ | $10.0 \pm 0.8$ |
| Mixed POPC-Cholesterol<br>(20%) | $7.6 \pm 0.8$ | $9.7 \pm 0.1$ |
| HAV-2B-POPC | $15.7 \pm 0.9$ | $20.4 \pm 0.4$ |
| HAV-2B-POPC<br>– Chol(20%) | $9.0 \pm 1.1$ | $11.2 \pm 0.7$ |

In order to understand the role of HAV-2B peptide in modulating lipid packing defects, we compare P(A) in presence of peptide with respect to that of each controls. We observe significant change in size distribution of both “Deep” and “Shallow” defects upon peptide insertion in comparison to pure POPC bilayer (Fig 5C). In POPC bilayer, the probability of finding “Deep” defects of size beyond 50 Å^2^ is trivial. However, the probability of finding similar sized defects not only increases but also occurrence of higher defect sizes ~ 80 Å^2^ become more probable under the influence of this peptide. Similar trend is observed for “Shallow” defects with presence of defect sizes upto 100 Å^2^ which are absent in the control system. This is reflected in increased area constants, Π, (see Table 1) of the two defect types. The “Deep” and “Shallow” defect area constants increase almost two-folds indicating that both the number of defect sites as well as the size of defects increase significantly upon peptide partitioning. Representative snapshots illustrating such enhanced prevalence and size of defect sites in HAV-2B-POPC system is shown in SI, Fig S8. On contrary, cholesterol embedded bilayer do not show much of a change in distribution of lipid packing defects, in presence or absence of the HAV-2B peptide (Fig 5D). The P(A) indicates prevalence of small defect sizes similar to control. As a result, the defect area constant (Table 1) is low and shows trivial change upon peptide adsorption. This is also evident upon comparison of representative snapshots of HAV-2B-POPC-Chol(20%) system (SI, Fig S8) with the respective control (Fig S7).

The presence of lipid packing defects can have profound implications in what the peptide, close to the membrane surface ‘sees’ in terms of transient binding sites before insertion and partitioning. We next focus on how the peptide can detect such difference in the lipid packing defects owing to different compositions of membrane and consequently how the partitioning is dictated.

#### (b) Hydrophobic insertion in co-localized defects

In order to understand the possible role of “Deep” packing defects in promoting HAV-2B peptide partitioning in POPC bilayer, we focus on the residues which undergo deep insertion into membrane milieu. These residues belonging to the C-terminal tail and the alpha-helical hairpin motif lie within ~ 10 Å of bilayer centre, quite deeply buried into the membrane. The insertion dynamics of these residues (see SI, Fig S6B) reveal a bulky hydrophobic residue, F56 partitions into the membrane as early as 160 ns, followed by other C-terminal residues and finally the hairpin motif. We scan the membrane topology to identify occurrence and location of “Deep” defects in vicinity of F56, preceding and succeeding the instance of such insertion event. Fig 6A shows the distance (in red) along z-direction of the centre of mass (C.O.M) of F56 relative to the average level of C2 atoms of glycerol moieties superposed on the identified “Deep” defect area (blue) overlapping with xy-coordinates of the inserting residue. Prior to insertion, we observe transient openings of “Deep” defects as F56 hovers close to the lipid-water interface. This instance is captured and illustrated through a representative snapshot at 147 ns (Fig 6B, top panel), where F56 is ~ 10 Å far from the level of glycerol while a “Deep” defect ~ 100 Å^2^ appears underneath. The defect vanishes with no insertion taking place, only to re-appear at frequent intervals, until at around 158 ns, a large defect opening of 150 Å^2^ facilitates quick approach of F56 (see mid panel of Fig 6B) towards the defect and subsequent insertion at 161 ns. The defect area at the point of insertion is 45 Å^2^, comparable to size of phenylalanine side chain [73]. Once inserted, we find an increase in the defect area at 166 ns so as to accommodate the bulky side chain of F56 resulting in deep partitioning into the membrane core (bottom panel Fig 6B). F56 remains inserted till the end of simulation (SI, Fig S6B), thus stabilizing this “Deep” defect.

**Fig 6.**
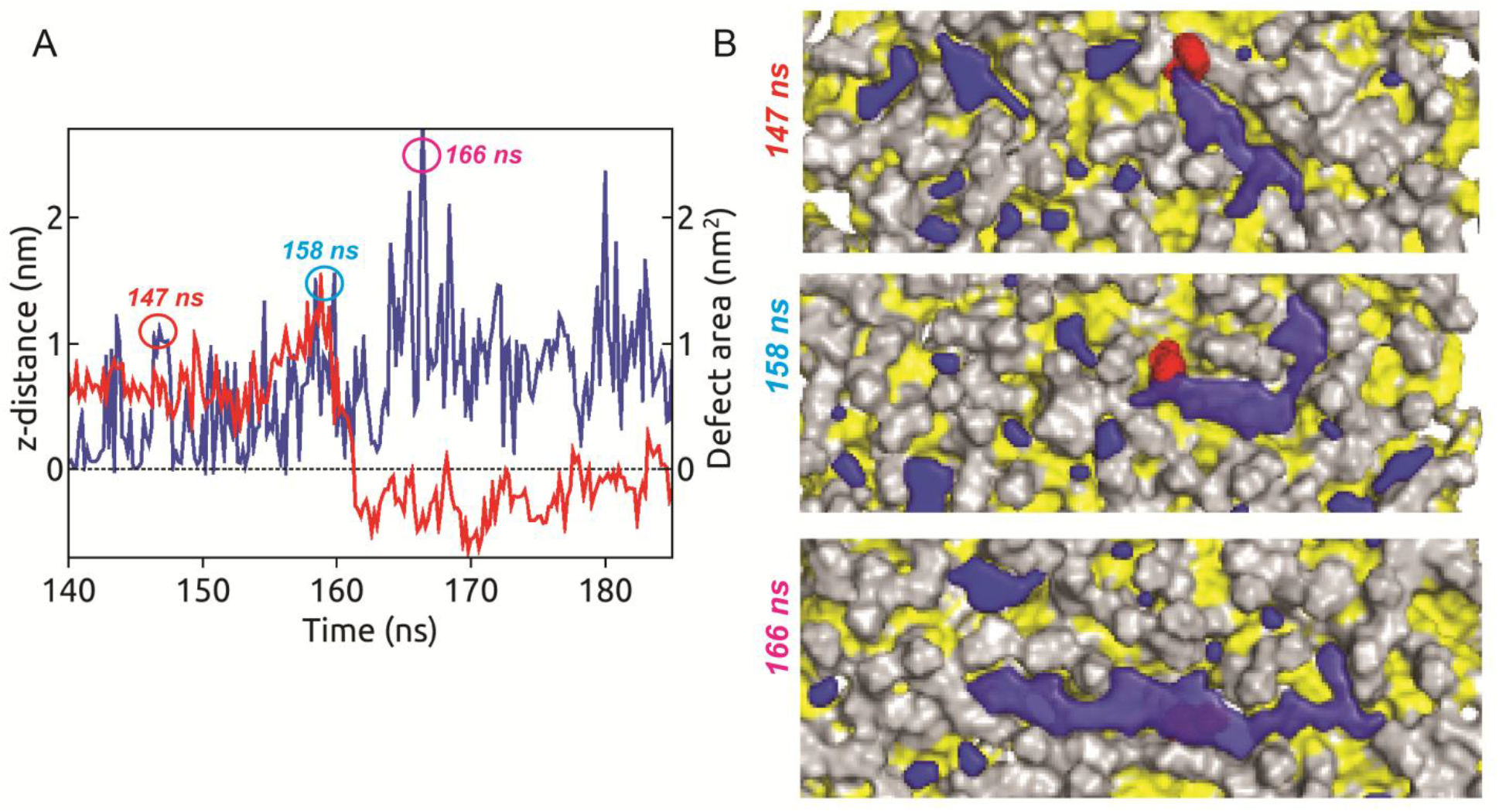
The co-localization of a large “Deep” packing defect and insertion of a bulky hydrophobic residue. (A) The time evolution of distance along z-direction (red) between center of mass (C.O.M) of F56 side-chain atoms and average coordinate of glycerol atoms superposed on the size of Deep packing defect (blue) co-localized on the same x-y coordinates as F56-C.O.M. The plot indicates instantaneous Deep defects opening in vicinity of F56 facilitating its sudden entry into membrane hydrophobic interior at around 160 ns. The insertion event is followed by an increase in defect area possibly to accommodate the bulky F56 side-chain. (B) A series of snapshots characterizing the sequence of insertion event are illustrated. The POPC headgroups (grey) and acyl chain (yellow) in surface representation are superposed with the co-localized “Deep” packing defect (in blue) along with F56 (in red).

We perform similar analysis to identify “Deep” defects co-localized with the hydrophobic residues of alpha helical hairpin motif. Although each of the hydrophobic residues undergoes discrete insertion events, interestingly, a single large co-localized defect is identified which facilitates deep partitioning of the entire hairpin motif. We illustrate the insertion dynamics of the entire hairpin motif considering the relative distance of its C.O.M from the average glycerol level along with the evolution of the co-localized “Deep” defect area in Fig 7A. We find the co-localized defect site extending upto 300 Å^2^ prior to complete insertion at around 338 ns. Subsequently, the defect is stabilized upon partitioning of the alpha helical hairpin motif, which remains embedded till the end. The final snapshot in Fig 7B shows complete insertion of the peptide into the large “Deep” defect site with the free N-terminal tail remaining solvent exposed.

**Fig 7.**
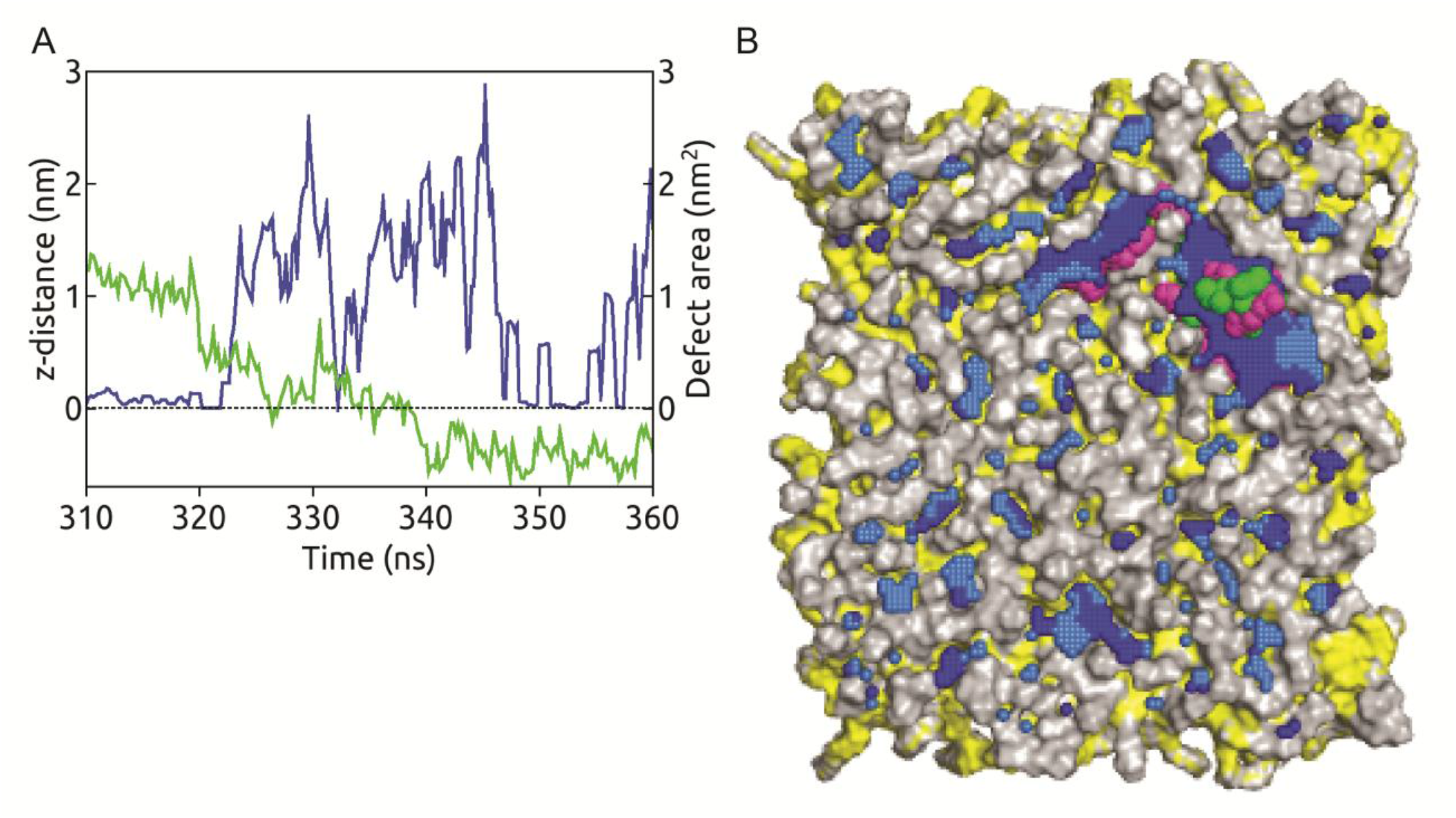
The co-localization of a large “Deep” packing defect and insertion of the alpha helical hairpin motif. The time evolution of distance along z-direction (green) between center of mass (C.O.M) of hydrophobic residues from alpha-helical hairpin motif and average coordinate of glycerol atoms superposed on the size of the co-localized “Deep” packing defect (blue). (B) Final snapshot of HAV-2B peptide inserted into the large co-localized “Deep” defect. The POPC headgroups (grey) and acyl chain (yellow) in surface representation are superposed with “Deep” (blue) and “Shallow” (light blue) packing defects along with the embedded peptide showing few exposed hydrophobic (green) and hydrophilic (magenta) residues.

We provide a comprehensive picture of mechanism of HAV-2B peptide insertion into the POPC membrane. The “Deep” defect sites pose as transient hotspots for hydrophobic insertion, promote deep partitioning of HAV-2B peptide into the membrane core and subsequently get stabilized upon peptide binding. The sequence of hydrophobic insertion, however, is expected to be stochastic. Owing to the dynamic nature of the C-terminal tail and presence of multiple hydrophobic residues with bulky side-chain, the discrete insertion events may be random depending on the availability of a nearby defect site. Nevertheless, the hydrophobic insertions being correlated to location of interfacial “Deep” defect sites indicate the ability of HAV-2B peptide in sensing lipid packing defects to gain entry into host membranes. On the contrary, there are fewer lipid packing defects in the control system when cholesterol is added. And it is this that prevents the partitioning of the peptide, because it is unable to find a binding hotspot / “hook” for hydrophobic insertion. This to our understanding provides the first insight into the entry mechanism of a non-enveloped viral lytic peptide into host membranes, which can be generalized to study other such systems.

## 3. DISCUSSSION

In the present work, we have shown that membrane remodelling due to change in bilayer composition or presence of active peptide regulate lipid packing defects. Early investigations by Voth and co-workers[70] revealed that lipid packing defects increased with an increase in membrane curvature, exhibiting curvature dependent characteristic defect area constant, Π (13.7 Å^2^ convex vs 7.8 Å^2^ flat). Later investigations confirmed a synergistic effect of membrane curvature and lipid tail unsaturation on interfacial packing defects.[79] Specifically, increasing the lipid acyl chain unsaturation or by introduction of conical lipids into flat bilayers, both size and number of defects increased considerably such that the defect size distribution of the flat bilayer resembled that of a positive curvature (convex) bilayer.[73, 78, 80] On the other hand, substitution of mono-unsaturated lipids with saturated ones led to scarcity of lipid packing defects. Our investigation on the role of cholesterol reveals quite a similar effect in modulating lipid packing defects. We observe a significant decrease in number of defect sites in presence of cholesterol. This may be understood based on the ordering and condensing effects of cholesterol. Owing to bilayer condensation, the lipid headgroups become more prone to aggregation. This is evident from the increased peak intensities of radial distribution function, g(r) of phosphate (P) atoms of lipid headgroups (SI, Fig S9) in presence of cholesterol. The enhanced aggregation of lipid headgroups and the ordering of lipid tails prevent solvent exposure of the hydrophobic tails, leading to reduced occurrence of defect sites.

Recently it was demonstrated that membrane thinning also enhances lipid packing defects. Based on MD simulations, Hilten et al[81] developed a protocol to vary membrane thickness by applying an external force. Upon comparing the defect distribution profile of POPC membrane with and without the thinning protocol, the authors reported higher probability of finding large defects in thin POPC membrane compared to a control of normal thickness. This is similar to our observations, where instead of the thinning protocol; the presence of the membrane active HAV-2B peptide in POPC induces global thinning accompanied by enhanced lipid packing defects. The induced thinning causes the POPC bilayer to expand laterally and consequently the P-P g(r) undergoes a decrease in peak intensity (SI, Fig S9) compared to the control system. The reduced aggregation propensity of POPC headgroups along with the enhanced disordering of its acyl chains facilitate frequent exposure of lipid tails to hydrating water, thus resulting in enrichment of defect sites in both number and size. As a consequence, the defect area constant (see Table 1) estimated from the defect size distribution profile of HAV-2B-POPC membrane is significantly higher compared to the control and resembles that of a convex bilayer[70].

In addition, the Hilten et al[81] study also showed that thinning induced enrichment of packing defects help sorting of lipids and protein. The authors reported preferential partitioning of cholesterol molecules in thick regions of the membrane away from the “defect-rich” thin regions, in contrast to localization of an amphipathic peptide near the thin zone of POPC membrane. On a similar note, we find that cholesterol-rich regions exhibit more thickness (Fig 3E-F). Further, a visual inspection of defect site snapshots of cholesterol based systems (Fig S7-S8) reveal partitioning of cholesterol molecules away from “Deep” defect sites but preferentially seem to localize close to “Shallow” defects. In the present work, we show that enrichment of “Deep” defects in presence of HAV-2B peptide facilitates hydrophobic insertion and deep partitioning into the membrane. We identify the presence of a single large “Deep” defect colocalized with the alpha helical hairpin motif that drives its partitioning and pronounced thinning around the insertion site. This provides a strong evidence of such thinness induced peptide sorting facilitated by defect sensing mechanism as demonstrated Hilten and co-workers[81].

Such partitioning mechanism being driven by sensing of lipid packing defects had been observed for hydrophobic nanoparticles[82], disaccharides[83] and other proteins[34, 71–76]. To this end, extensive studies had been performed on amphipathic lipid packing sensor (ALPS) motif peptide[34, 71, 73, 81]. The ALPS motif commonly found in several peripheral membrane binding proteins, is a short (20-40 amino acid) amphipathic alpha-helix. The amino acid distribution is quite similar to our peptide, being rich in polar and bulky hydrophobic residues but lacking (or few) charged ones. Vanni et al[73] demonstrated that the ALPS motif partitioning progresses via detection of lipid packing defects and subsequent insertion of bulky hydrophobic residues into large pre-existing “Deep” defects. The authors observed that ALPS motif insertion caused minor modifications in the distribution of “free” packing defects compared to that of the peptide-free control bilayer and concluded that no new defects are formed under the influence of the peptide. However, recently, it had been shown that new defect formation or enhancement of a packing defect size occur with the ALPS motif in close proximity to membrane and that preexistence of a large packing defect is not necessary for peptide insertion.[71] We also check if a similar effect of HAV-2B peptide exists on lipid packing defects in the following paragraph.

To this end, we consider the action of HAV-2B peptide on lipid packing “Deep” defects at pre- and post-insertion time-scales barring the first 50 ns for top and bottom leaflets separately. The time evolution of defect area fraction, *f* and number of defect sites, *n_sites_* are shown in Fig S10A and B respectively. It is important to note that even prior to insertion of the alpha helical hairpin motif at around 338 ns, the defect area fraction (Fig S10A) is significantly higher compared to the POPC control (see Fig 5A) and is also similar in both leaflets, although the peptide is more close to the top than the bottom leaflet. After the insertion event, however, the defect area fraction of top leaflet increases more due to stabilization of the large “Deep” defect that harbours the peptide. The *nsites* (Fig S10B) for both leaflets remain stable and similar throughout but considerably higher than the control (see Fig 5B). We generate the size distribution of defects prior to peptide insertion between 50 to 250 ns and compare with the post-insertion (250-500 ns) distribution (see Fig S10C). The post-insertion distribution profile of “Deep” defect size in HAV-2B-POPC system shown in Fig S10C is same as that in Fig 5C, except for the fact that the distribution is truncated to data fitting range in Fig 5C. Both the pre- and post-insertion size distribution profiles are overlapping for moderately large defects upto ~ 100 Å^2^ and follow single exponential decay with similar area constants ~ 16 Å^2^. The comparison of pre- and post-insertion distributions of such defect sizes clearly indicate that mere presence of the peptide close to the bilayer induces formation of new defect sites as well as enlargement of defect sizes in both leaflets compared to control POPC bilayer. This effect of HAV-2B peptide may be attributed to its membrane active property, unlike the ALPS motif, which, being associated with peripheral membrane proteins lack such activity. Such action of HAV-2B peptide hints at a co-operative process involving membrane re-modelling that enhances lipid packing defects favouring discrete hydrophobic insertion events, which in turn stabilizes the packing defect leading to destabilization / remodelling of membrane.

Significant differences in the pre- and post-insertion distributions are observed for defect sizes beyond 100 Å^2^. Moreover deviation from single exponential behaviour is also observed in this regime, consistent with earlier observations [70, 71]. The post-insertion distribution is bimodal with a second peak appearing close to 300 Å^2^ due to stabilization of the large “Deep” defect upon partitioning of the alpha helical hairpin motif. Since formation of large defects become more probable in presence of the HAV-2B peptide, we track formation of the largest (maximum) defect size in each frame and in each leaflet separately and plot the time evolution of the same (see Fig S11). The time-profile of maximum defect size in the top leaflet show characteristic spikes before the complete insertion of the alpha helical hairpin motif. The spikes indicate momentary openings of very large defects ranging between 100-200 Å^2^ between 50-300 ns and thus explain the deviation from single exponential behaviour observed for large defect sizes. But these spikes are conspicuously absent in the lower leaflet which shows stable values (~ 60 Å^2^) of maximum defect size openings. We have shown earlier in the Results section corresponding to the hydrophobic insertion events (Figs 6–7) that co-localized defects show increase in defect area as the distance between hydrophobic residue and the defect decreases. Hence, the approach of HAV-2B peptide towards the upper leaflet elicits such momentary large defect site openings giving rise to the observed spikes, consistent with that observed for ALPS binding[71]. These defect openings often correlate with discrete insertions of bulky hydrophobic residues but not always as indicated in Fig S11. If the peptide is unfavourably oriented insertion may not proceed, like the first two spikes do not correspond to any binding events. Finally, just prior to insertion of the alpha helical hairpin, still larger defect openings > 200 Å^2^ is observed close to 340 ns, until a large defect ~ 300 Å^2^ gets stabilized by its insertion. In comparison, the ALPS binding reported large defect size ~ 115 Å^2^.[71] Owing to the hairpin structure of HAV-2B with two helices, the stabilized “Deep” defect is comparatively larger than the one observed for ALPS binding. These results strongly suggest that presence of lipid packing defects, which are a function of membrane composition, act as transient albeit weak binding spots for many membrane active agents in the initial phase of recognition and binding prior to membrane partitioning. Our results also show how a representative of a class of viroporins may exploit such packing defects for deep insertion into host cell membranes, a requisite step in viral entry mechanism. Whether similar mechanism is adopted by other viroporin families is a part of future studies.

## 4. CONCLUSIONS

In conclusion, we provide a comprehensive picture of mechanism of HAV-2B peptide insertion and partitioning into POPC membrane, and thereby causing membrane disruption in agreement to experimental studies. Our simulations strongly suggest that the peptide is capable of sensing lipid packing defects which facilitates peptide partitioning dynamics through hydrophobic residue insertions into membrane milieu. The peptide insertion further destabilizes the membrane by inducing global thinning and enhanced disordering of lipid tails. We also show that the presence of cholesterol significantly alters such lipid packing defects so as to mitigate partitioning into cholesterol-rich membranes. Our current study provides first atomistic insight into the entry mechanism of a non-enveloped viral lytic peptide into host membranes via lipid packing defects, and can possibly be a part of general membrane detection mechanism for the viroporin class of viruses.

## 5. METHODS

Since, the structure of membrane active part of HAV-2B peptide is yet to be determined experimentally, it has been modelled in a recent study [10] and we use this modelled peptide in our simulation study. It is 60 amino acids long (see primary sequence in SI, Fig S1) and typically unstructured with an α-helical hairpin motif possessing viroporin like membrane permeation activity. The protonation states of peptide residues are calculated using PROPKA3 [84, 85] at neutral pH and hydrogen atoms are added accordingly. The peptide is then solvated in a water box generated by considering a thickness of 10 Å from peptide surface followed by neutralization with addition of salt ions to maintain a concentration of 0.15 M. A pure 1-palmitoyl-2-oleoyl-sn-glycero-3-Phosphatidylcholine (POPC) bilayer is generated using CHARMM-GUI Membrane Builder [86–88]. The bilayer is built in a rectangular box with surface area of 100×100 Å^2^, comprising of 147 POPC molecules per leaflet and hydration number ~ 35 water molecules per lipid, representing a fully hydrated fluid phase as reported from experiments [89]. Similarly, with 117 POPC and 30 cholesterol molecules per leaflet, representing a mixed bilayer of POPC-Cholesterol (20%) is prepared. Further, required counterions are included to achieve a salt concentration of 0.15 M. We take the peptide and the bilayers, after proper equilibration of individual systems, to generate a system representing peptide at lipid-water interface to understand effects of HAV-2B on membrane systems with and without cholesterol. The peptide is placed with its centre of mass at a distance of 15 Å from the upper leaflet of the bilayers avoiding any steric clashes, extra water molecules and ions are added to ensure proper solvation and desired salt concentration.

Molecular dynamics (MD) simulations for each of these systems (see SI Table S1) are performed in NAMD2.10 [90] using modified TIP3P [91] water model, CHARMM36m [92] and CHARMM36 [93] force field parameters for the peptide and the lipid molecules, respectively. All systems are energy minimized for 10,000 steps followed by equilibration by gradually turning off restraints. The simulations are carried out using periodic boundary conditions in NPT ensemble at 1 atm pressure and 310K with a time step of 2 femto-seconds. The van der Waals interactions are smoothly truncated beyond 12 Å, by a forced-based switching function between 10 Å and 12 Å, while Particle mesh Ewald fast Fourier transform is used for electrostatic interactions. The total simulation time and equilibration time of each system is indicated in Table S1. The equilibrated trajectories are used for further analysis using Visual Molecular Dynamics (VMD) [51], MEMBPLUGIN [94], Packmem[78] and in-house Fortran codes.

## AUTHOR CONTRIBUTIONS

MB and SV designed the project. SS performed the simulations and carried out the analysis. All authors contributed to writing and reviewing the manuscript.

## ACKNOWLEDGEMENT

All simulations in this work have been carried out on supercomputing facility Nandadevi cluster at The Institute of Mathematical Sciences, Chennai, India.

