## Supplementary Information for "Effect of Cholesterol on Membrane Partitioning Dynamics of Hepatitis A Virus-2B peptide"

| Systems | System size | Simulation time (ns) | Equilibration time (ns) |
|---|---|---|---|
| HAV-2B – Water | 22288 | 400 | 100 |
| Pure POPC | 70306 | 350 | 150 |
| Mixed POPC – Chol(20%) | 66710 | 350 | 150 |
| HAV-2B – POPC | 104280 | 500 | 300 |
| HAV-2B – POPC - Chol(20%) | 93379 | 715 | 450 |

Table S1. Details of systems considered in the present study. All atom MD simulations of HAV-2B are performed in different media: Water, hydrated POPC bilayer, hydrated POPC-Chol(20%) bilayer. Pure POPC and Mixed POPC-Chol(20%) refers to control membrane systems in absence of the peptide. The system size denoted by total number of atoms; followed by the simulation time and the equilibration time for each system are provided as well.

| *Residue id* | *HAV-2B - POPC* | *HAV-2B – POPC-Chol(20%)* | *Residue id* | *HAV-2B - POPC* | *HAV-2B – POPC-Chol(20%)* | *Residue id* | *HAV-2B - POPC* | *HAV-2B – POPC-Chol(20%)* |
|---|---|---|---|---|---|---|---|---|
| V1 | 36.9 | 0 | V21 | 100.0 | 0 | N41 | 100.0 | 13.4 |
| T2 | 11.4 | 0 | I22 | 100.0 | 0 | Y42 | 100.0 | 12.5 |
| V3 | 0 | 0 | Q23 | 100.0 | 0 | A43 | 100.0 | 0 |
| E4 | 0 | 0 | Q24 | 100.0 | 0 | D44 | 94.5 | 2.6 |
| I5 | 0 | 0 | L25 | 100.0 | 0 | I45 | 100.0 | 0 |
| I6 | 0 | 0 | N26 | 100.0 | 0 | G46 | 97.2 | 0.9 |
| N7 | 0 | 0 | Q27 | 100.0 | 0 | C47 | 100.0 | 2.6 |
| T8 | 0 | 0 | D28 | 99.2 | 0 | S48 | 100.0 | 53.5 |
| V9 | 0.4 | 0 | E29 | 62.3 | 0 | V49 | 100.0 | 58.7 |
| L10 | 8.6 | 0 | H30 | 83.5 | 0 | I50 | 100.0 | 100.0 |
| C11 | 7.8 | 0.29 | S31 | 49.4 | 0 | S51 | 100.0 | 91.3 |
| F12 | 91.0 | 7.8 | H32 | 89.8 | 0 | C52 | 94.5 | 24.1 |
| V13 | 32.2 | 1.4 | I33 | 100.0 | 0 | G53 | 100.0 | 41.9 |
| K14 | 93.7 | 46.2 | I34 | 97.6 | 0.6 | K54 | 100.0 | 79.6 |
| S15 | 100.0 | 54.9 | G35 | 100.0 | 0 | V55 | 100.0 | 13.7 |
| G16 | 100.0 | 9.3 | L36 | 100.0 | 0 | F56 | 100.0 | 5.8 |
| I17 | 100.0 | 1.4 | L37 | 100.0 | 0 | S57 | 100.0 | 10.2 |
| L18 | 100.0 | 0 | R38 | 99.2 | 6.7 | K58 | 99.2 | 1.2 |
| L19 | 100.0 | 0 | V39 | 100.0 | 0 | M59 | 100.0 | 1.2 |
| Y20 | 100.0 | 0 | M40 | 100.0 | 0 | L60 | 100.0 | 1.2 |

Table S2. Insertion probability of HAV-2B peptide residues into membrane computed over last 50 ns of equilibrated trajectories. The residues showing insertion probability > 75% are highlighted in colour according to its nature: Hydrophobic (Green) and Hydrophilic (magenta).

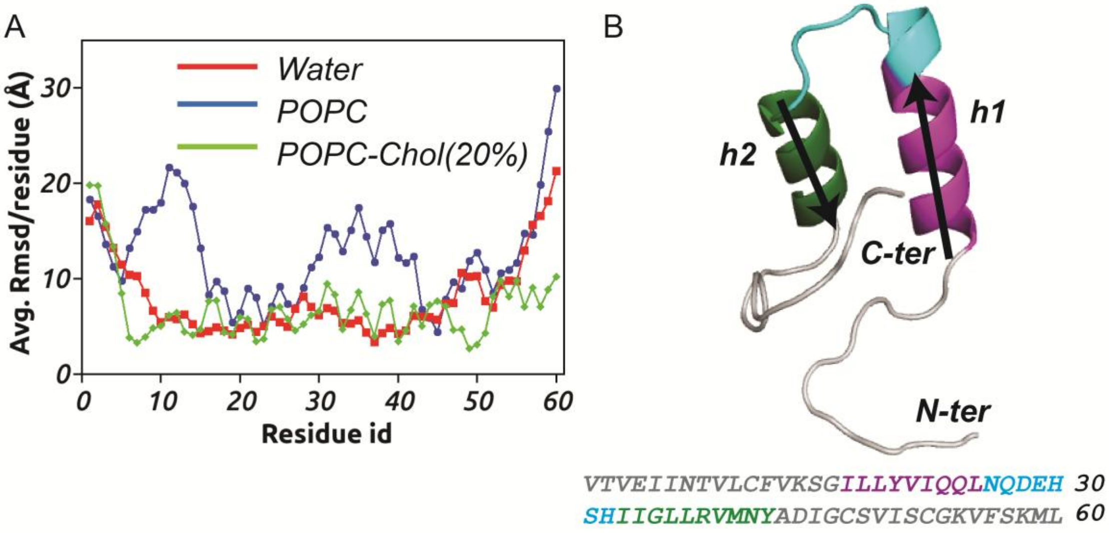


Fig S1. (A) Average RMSD per residue of HAV-2B peptide in water (red), POPC-water interface (blue) and POPC-Chol(20%)-water interface (green). (B) The average structure of HAV-2B peptide in water showing the unstructured N- and C-termini tails (in gray) and the structured hairpin motif characterized by presence of two alpha-helices (in magenta and green) connected by a turn (cyan). The same colours have been used to show the amino acid sequence of the peptide. The two helical axes are indicated as ***h1*** and ***h2***, used for calculation of inter-helical crossing angle, Ω.

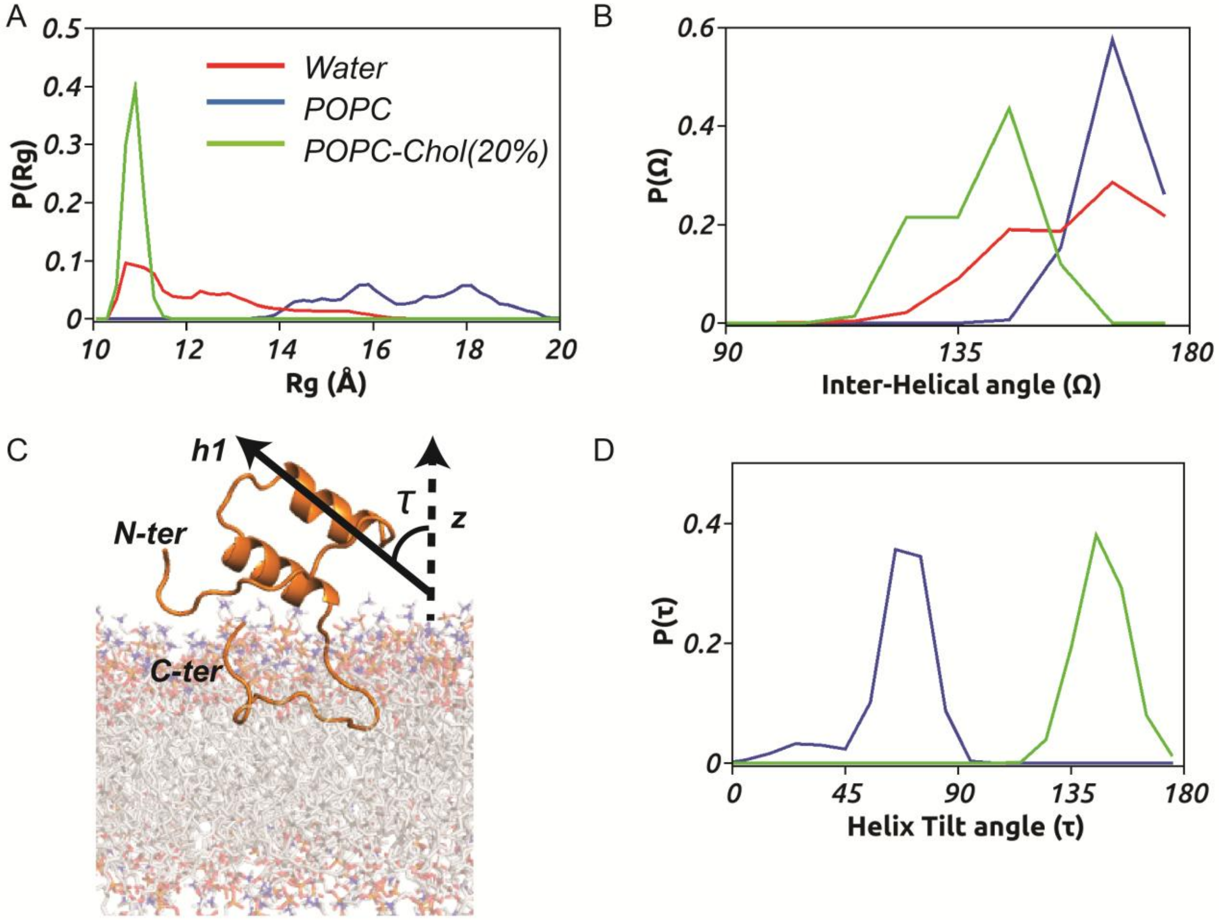


Fig S2 (A) Probability distribution of radius of gyration, P(Rg) of HAV-2B peptide in water (red), POPC (blue) and POPC-Chol(20%) bilayers (green). (B) Probability distribution of inter-helical crossing angle, P(Ω) of HAV-2B peptide in water (red), POPC (blue) and POPC-Chol(20%) bilayers (green). (C) A representative snapshot of peptide-membrane system indicating the helix tilt angle, τ of first helix axis (***h1***) with respect to bilayer normal, ***z*** and (D) The corresponding probability distribution, P(τ) of HAV-2B peptide in POPC (blue) and POPC-Chol(20%) bilayers (green).

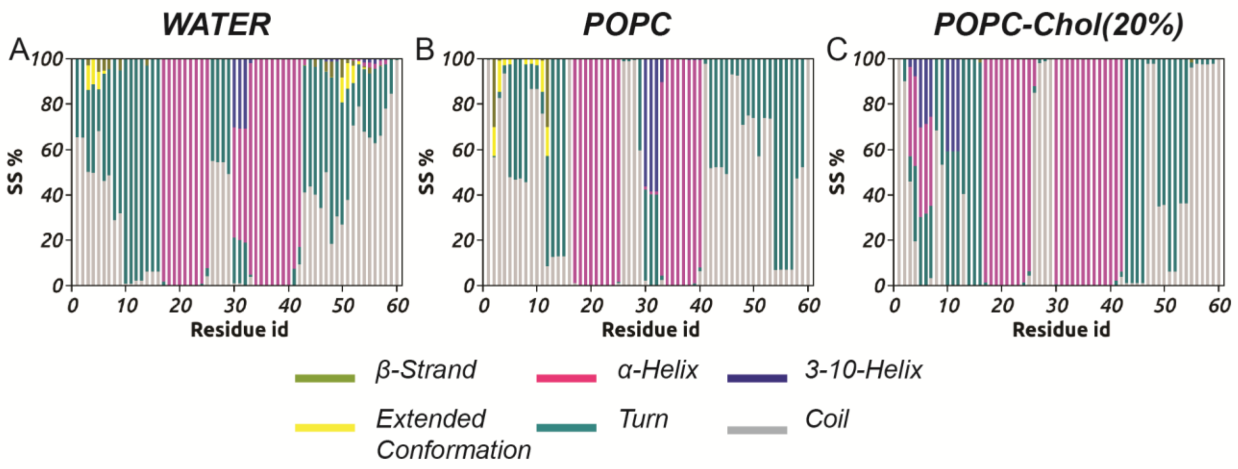


Fig S3. Residue based secondary structure (SS) % of HAV-2B peptide in (A) water, (B) in POPC and (C) in POPC-Chol(20%) bilayers, showing population of different secondary structural elements accessible to each residue.

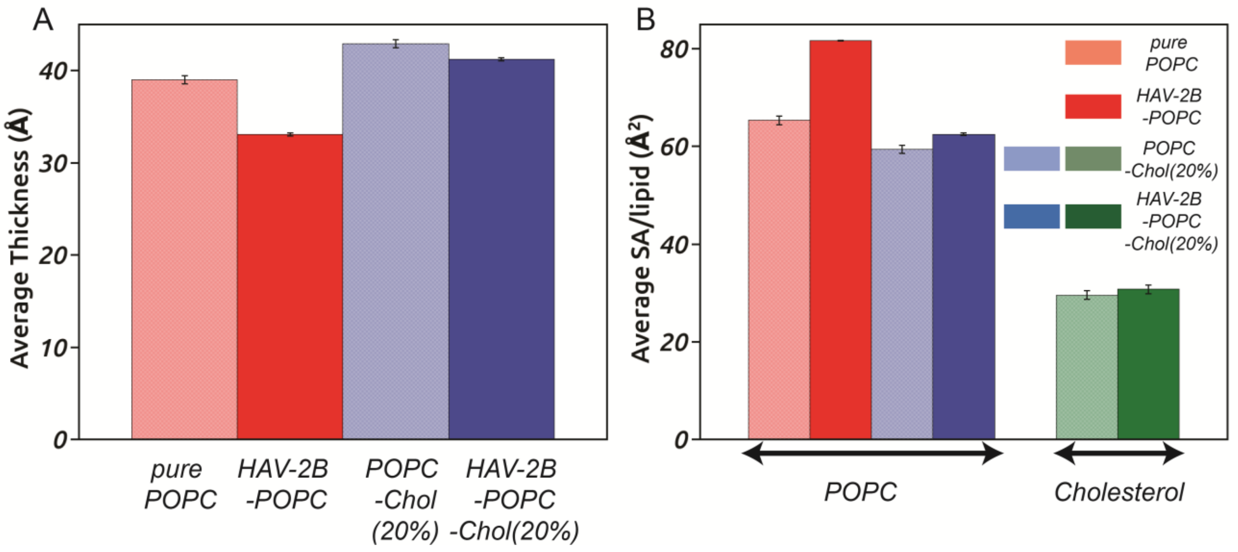


Fig S4. (A) Average bilayer thickness and (B) average surface area (SA) per lipid for different systems, computed over equilibrated trajectories.

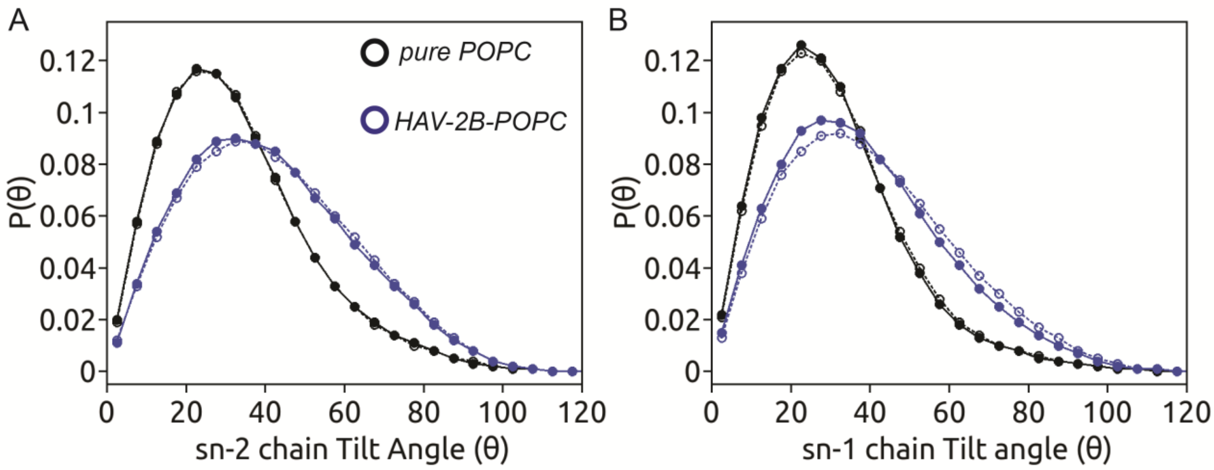


Fig S5. The distribution of lipid tail tilt angles for (A) sn-2 and (B) sn-1 chains in control pure POPC (black) and HAV-2B-POPC (blue) bilayers. The values for top leaflet are shown in solid symbol and solid line, while that of bottom leaflet are shown in open symbol and dotted line.

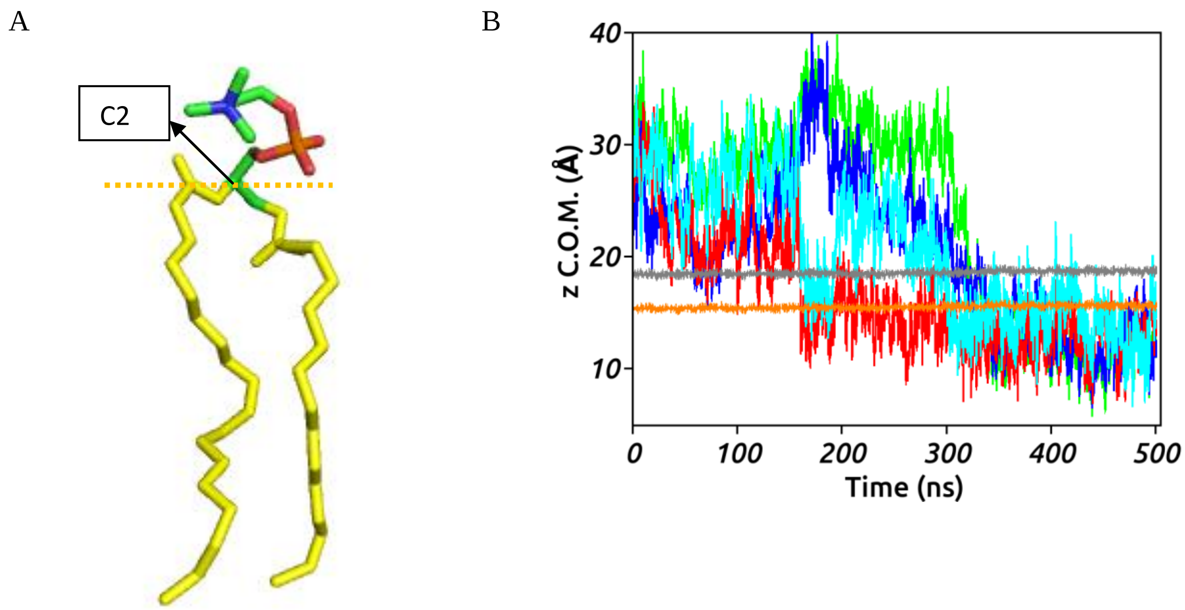


Fig S6 (A). C2 atom of glycerol moiety of POPC molecule based on the level of which defects are classified as “Deep” or “Shallow” . (B) The insertion dynamics of M40 (green), Y42 (blue), F56 (red) and M59 (cyan) residues from C-terminal tail of HAV-2B peptide, which show maximum insertion depth in

POPC bilayer. The average levels of POPC haedgroup atoms, P (grey) and glycerol, C2 (orange) are shown. Insertion of F56 occurs at 160 ns and remains embedded below the C2 level for rest of the time.

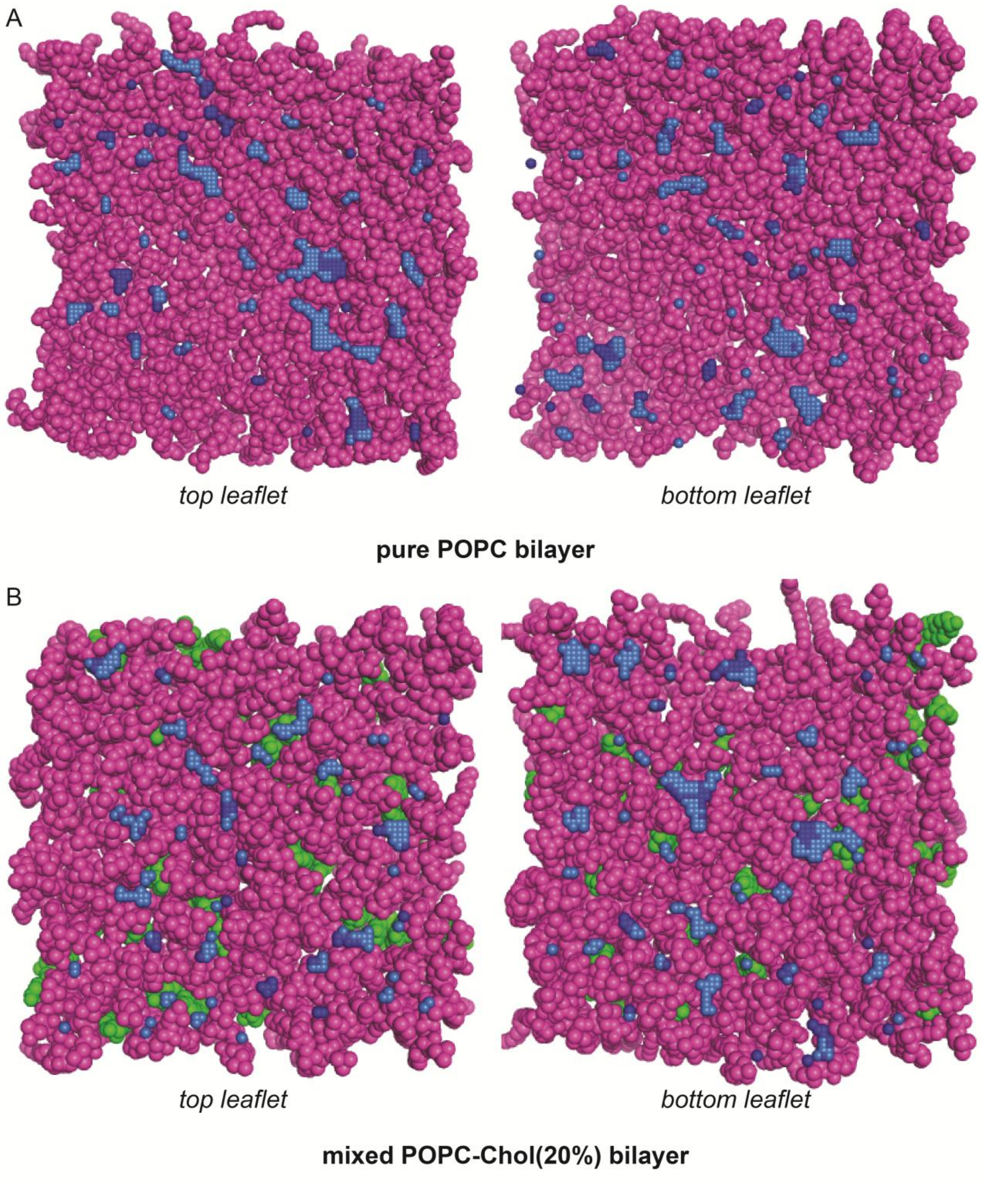


Fig S7. Representative snapshots of final MD structures showing packing defects in each leaflet of the control systems. POPC (magenta) and cholesterol (green) along with superposed "Deep" (dark blue) and "Shallow" (light blue) lipid packing defects are shown.

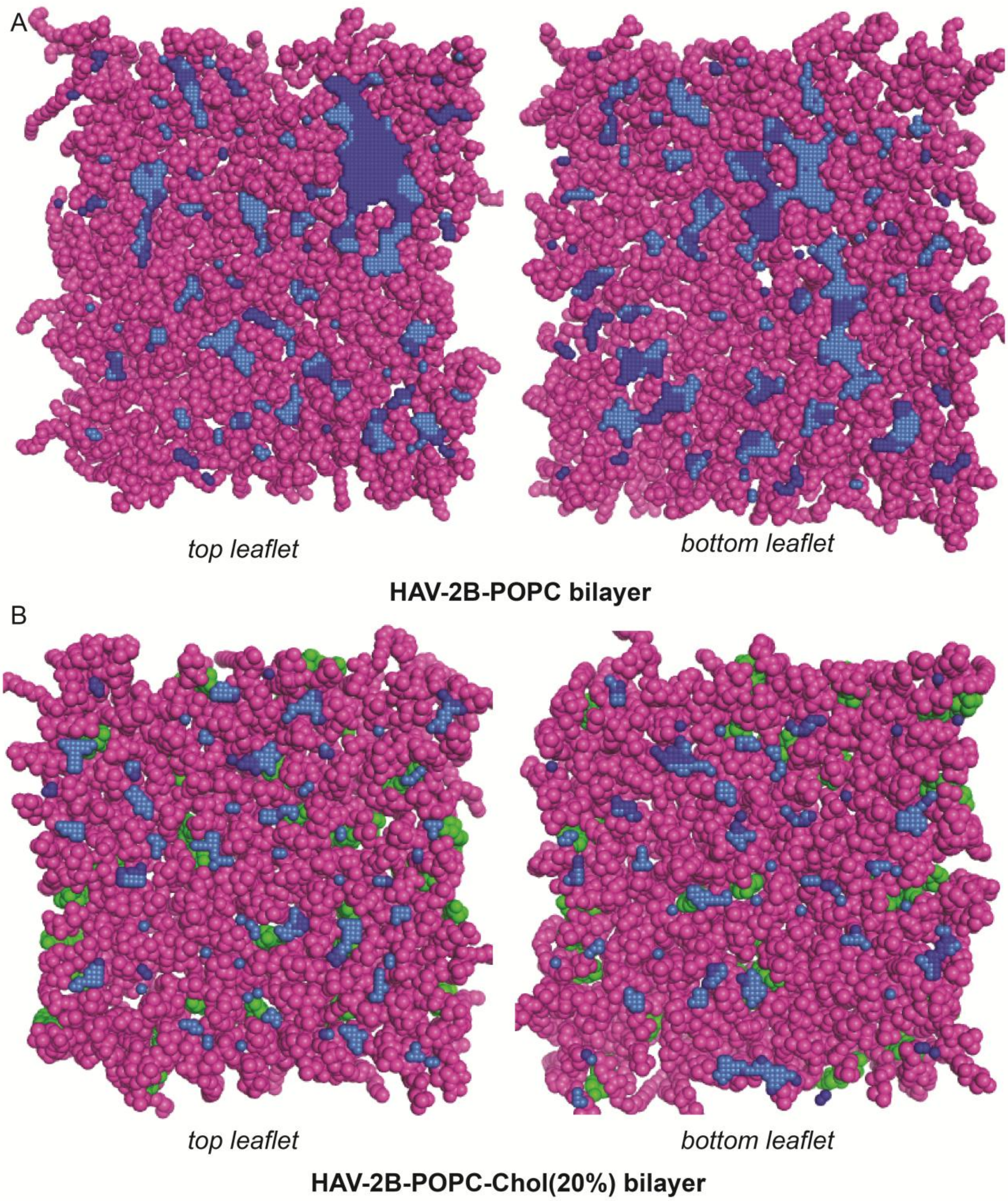


Fig S8. Representative snapshots of final MD structures showing packing defects in each leaflet of the peptide-membrane systems. POPC (magenta) and cholesterol (green) along with superposed “Deep” (dark blue) and “Shallow” (light blue) lipid packing defects are shown.

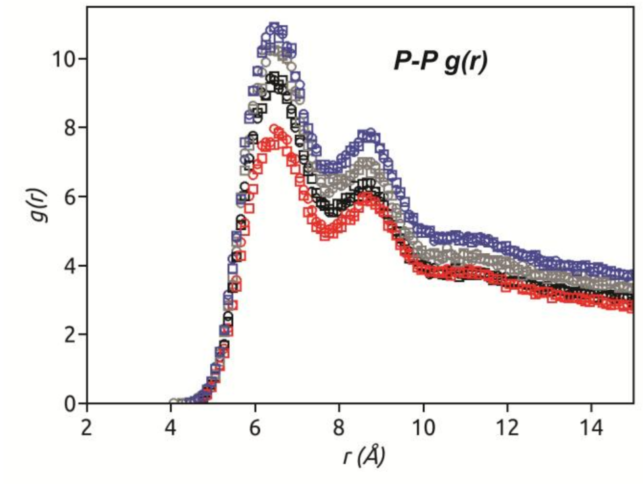


Fig S9. Radial distribution function of POPC headgroups considering phosphate (P) atoms of top (circle) and bottom (square) leaflets of different systems: pure POPC (black) and in presence of HAV-2B peptide (red), mixed POPC-Cholesterol (20%) (gray) and in presence of peptide (blue).

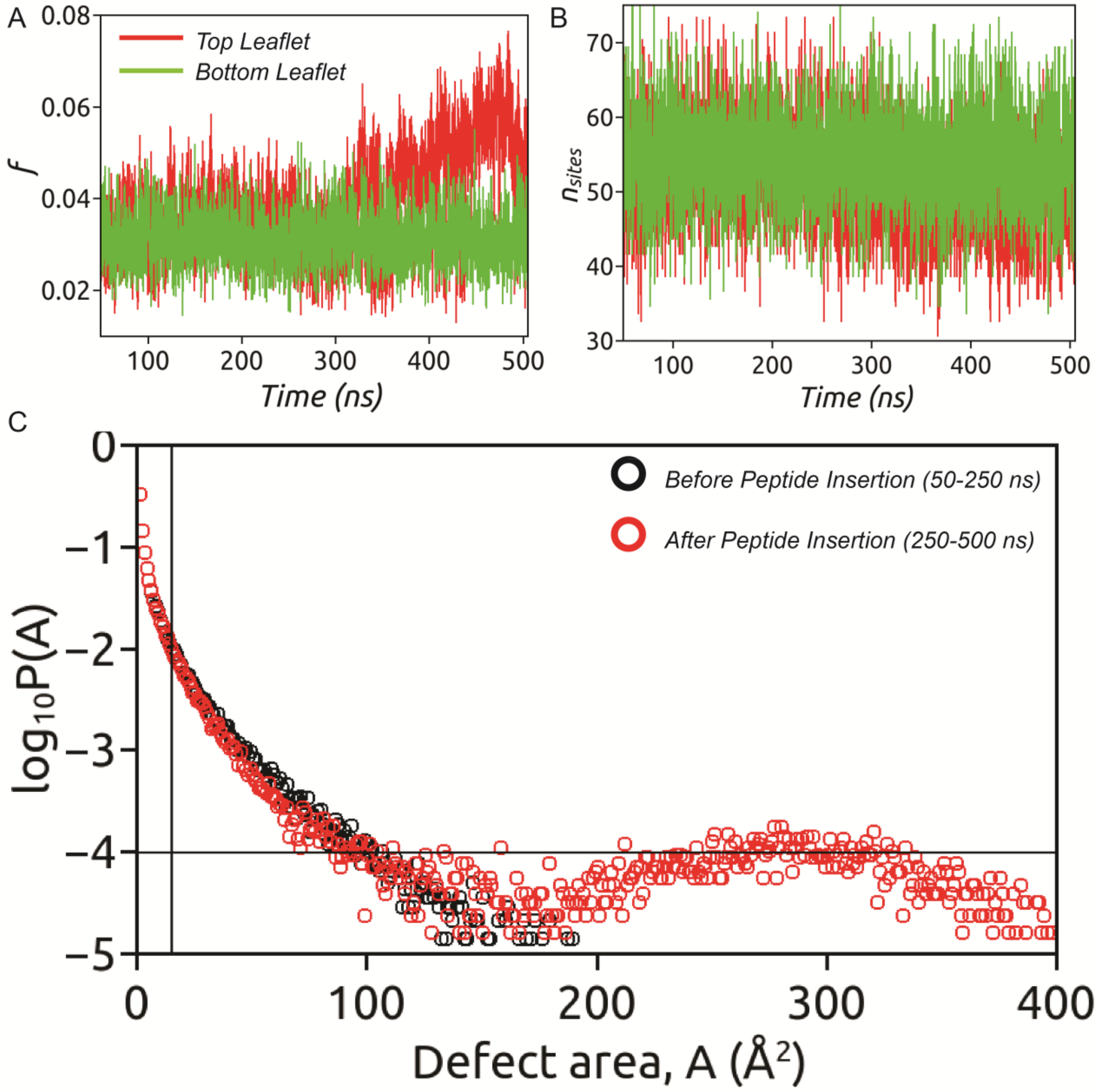


Fig S10. The effect of HAV-2B peptide partitioning on (A) "Deep" defect area fraction, $f$ and (B) number of "Deep" defect sites per leaflet ($n_{sites}$), as a function of time. (C) The probability distributions of defect sizes (in Å$^2$), $\log_{10}P(A)$, computed separately for 50-250 ns (before peptide insertion) and 250-500 ns (after peptide insertion).

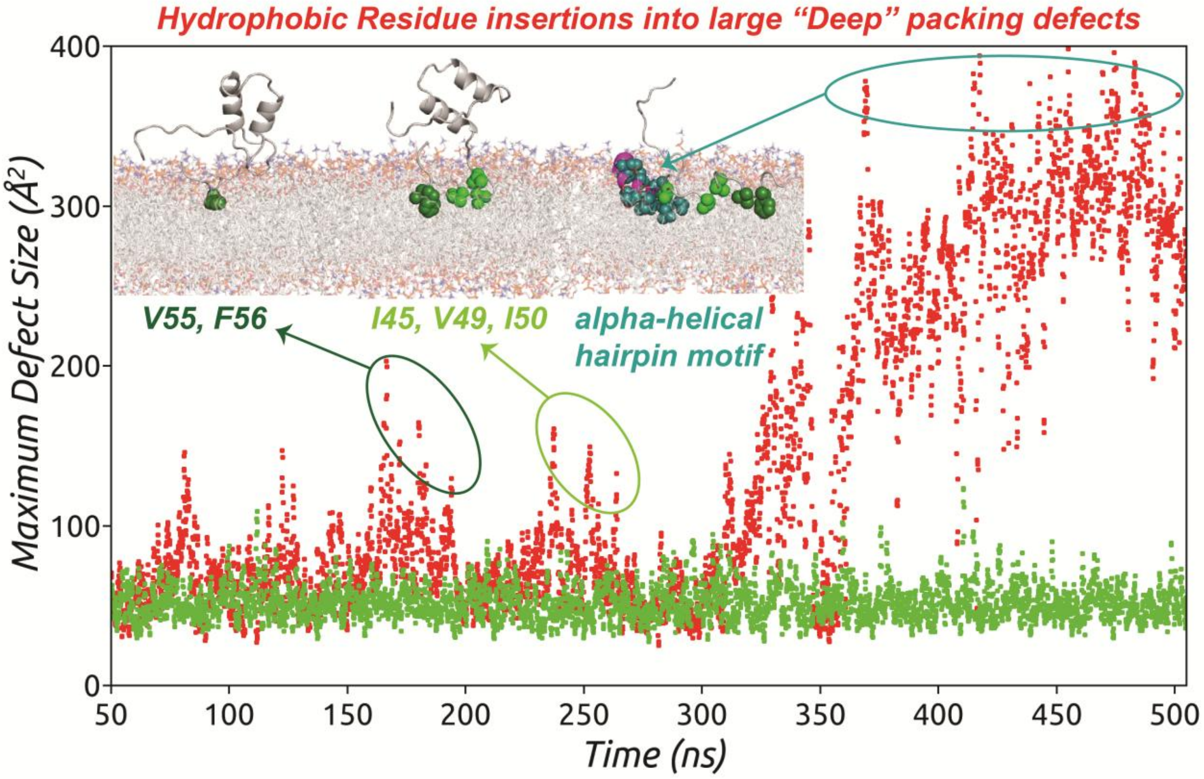


Fig S11. The time evolution of maximum "Deep" defect size occurring in top (red) and bottom (green) leaflets under the influence of HAV-2B peptide. Spikes appearing in maximum "Deep" defect size of top leaflet correspond to discrete hydrophobic residue insertion events. The complete insertion of the amphipathic alpha helical hairpin motif stabilizes a very large "Deep" defect of area > 300 $Å^2$.
